# Flow-induced mechanical coupling between perinuclear actin cap and nucleus governs spatiotemporal regulation of YAP transport

**DOI:** 10.1101/2022.11.15.516697

**Authors:** Tianxiang Ma, Xiao Liu, Haoran Su, Yuan He, Fan Wu, Chenxing Gao, Kexin Li, Zhuqing Liang, Dongrui Zhang, Xing Zhang, Ke Hu, Shangyu Li, Li Wang, Min Wang, Shuhua Yue, Weili Hong, Xun Chen, Jing Zhang, Xiaoyan Deng, Pu Wang, Yubo Fan

**Affiliations:** Beijing Advanced Innovation Centre for Biomedical Engineering, Key Laboratory for Biomechanics and Mechanobiology of Chinese Education Ministry, School of Biological Science and Medical Engineering, Beihang University, Beijing 100083, China; School of Engineering Medicine, Beihang University, Beijing 100083, China; Biomedical Pioneering Innovation Center (BIOPIC), Peking University, Beijing 100871, China; Academy for Advanced Interdisciplinary Studies, Peking University, Beijing 100871, China; Department of Gynecology and Obstetrics, Strategic Support Force Medical Center, Beijing 100101, China

**Author notes:** These authors contributed equally to this work (Tianxiang Ma, Xiao Liu, Haoran Su). Address for correspondence: Dr. Liu, Dr. Wang and Dr. Fan, School of Biological Science & Medical Engineering, Beihang University, Beijing 100191, China.

**Keywords:** Flow shear stress, YAP, Cellular mechanotransduction, Nucleocytoplasmic transport, Actin cap, Mechanochemical modeling

## Abstract

Mechanical forces, including flow shear stress, regulate fundamental cellular process by modulating the nucleocytoplasmic transport of transcription factors, such as Yes-associated Protein (YAP). However, the mechanical mechanism how flow induces the nucleocytoplasmic transport remains largely unclear. Here we found that unidirectional flow applied to endothelial cells induces biphasic YAP nucleocytoplasmic transport with initial nuclear import, followed by nuclear export as perinuclear actin cap forms and nuclear stiffening in a dose and timing-dependent manner. In contrast, pathological oscillatory flow induces slight actin cap formation and nuclear softening, sustaining YAP nuclear localization. To explain the disparately spatiotemporal distribution of YAP, we developed a three-dimensional mechanochemical model considering coupling processes of flow sensing, cytoskeleton organization, nucleus mechanotransduction, and YAP spatiotemporal transport. We discovered that actin cap formation and nuclear stiffness alteration under flow synergically regulate nuclear deformation, hence governing YAP transport. Furthermore, we expanded our single cell model to a collective vertex framework and found that actin cap irregularities in individual cells under flow shear stress potentially induce topological defects and spatially heterogeneous YAP distribution in cellular monolayers. Our work unveils the unified mechanism of flow-induced nucleocytoplasmic transport, offering a universal linkage between transcriptional regulation and mechanical stimulation.

---

Living cells are constantly subjected to distinct mechanical cues, such as matrix viscoelasticity^1^ and compressive forces^2^, and flow shear stress^3–5^. These forces can transmit to the nucleus through the cytoskeleton, causing nuclear deformation and regulating the nucleocytoplasmic transport of transcription factors^6–8^, such as Yes-associated Protein (YAP)^9, 10^. Flow shear stress is particularly relevant for cells in contact with fluids like blood, lymphatic, and interstitial flow, profoundly influencing cellular processes and diseases, such as atherosclerosis and cancer^3–5^. Although chemical pathways governing YAP’s spatiotemporal distribution have been identified under various flow conditions^5, 11–13^, the mechanical mechanism how flow induces the nucleocytoplasmic transport remains largely unclear.

Flow shear stress can induce perinuclear actin cap formation^14^, which tightly connects to the nucleus and exerts higher stress than conventional fibers^15^. Additionally, nuclear stiffness itself affects nuclear deformation^8, 16^. Based on these clues, we hypothesize that the YAP nucleocytoplasmic shuttling under different flow conditions is regulated by the nuclear deformability, which is controlled by the actin reorganization and nuclear mechanics alteration. We combined microfluidic vascular chip and Brillouin microscopy, and found that unidirectional flow induces biphasic YAP nucleocytoplasmic transport with initial YAP nuclear import, followed by nuclear export as actin cap formation and nuclear stiffening occur. In contrast, oscillatory flow maintains YAP nuclear localization through slight actin cap formation and nuclear softening. We then developed a mechanochemical model accounting for flow sensing at the plasma membrane, subsequent effects on the cytoskeleton, force transmission to the nucleus, and impact on YAP transport. Our findings reveal that actin cap concentrates stress on linkers of nucleus to cytoskeleton (LINC) complex while reducing overall nuclear membrane stress. The increasing nuclear stiffness collaboratively inhibit nuclear membrane deformation, ultimately governing the nucleocytoplasmic transport of YAP. Furthermore, we extend this single-cell framework – in a minimal way – to the analysis of cell monolayer, and predict that irregularities in actin cap formation within individual cells dictate the collective topological defects and spatially heterogeneous YAP distribution in monolayer.

## Flow-induced actin cap formation and nuclear stiffness alterations are associated with YAP nucleocytoplasmic transport

Expanding on previous investigations of endothelial cell morphological response to flow^17^, we examined the distinct effects of unidirectional and oscillatory shear stress on actin cap formation, nuclear stiffness, and YAP nucleocytoplasmic distribution using a microfluidic system (Extended Data Fig. 1a-c).

**Fig. 1.**
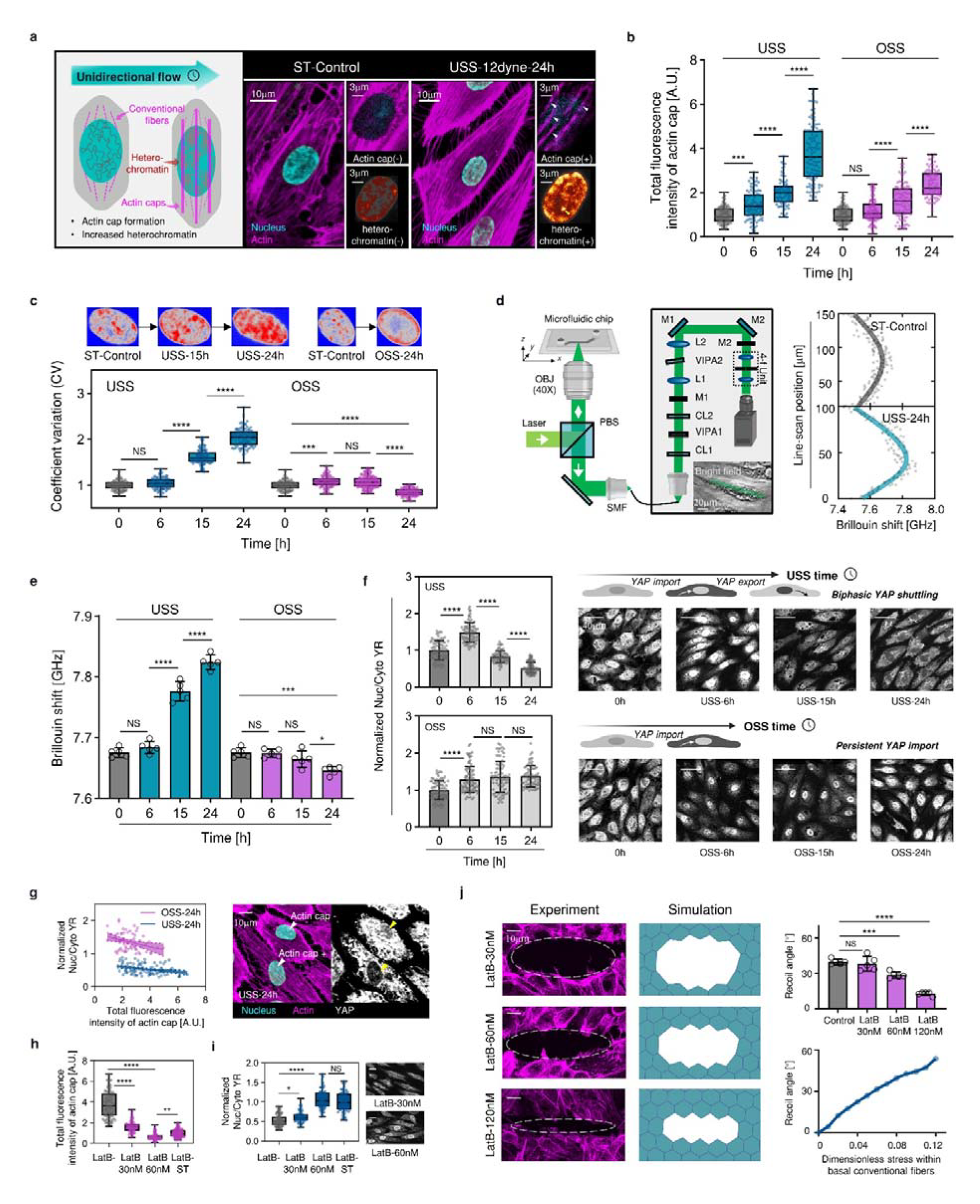
The connection between actin cap formation, nuclear mechanics alterations, and YAP localization under flow shear stress. **a**, Schematic of endothelial cell subjected to unidirectional shear stress with increasing actin cap formation and heterochromatin (stained by DAPI and phalloidin for nucleus and actin filament). Arrowhead indicates the actin cap and nuclear heterochromatin. **b,c**, Total fluorescence intensity of actin cap and coefficient of variation (CV) in response to unidirectional shear stress (USS, 12dyne/cm^2^) and oscillatory shear stress (OSS, 0±12dyne/cm^2^, 1Hz) (3 independent experiments, n > 80 cells). **d**, Setup of label-/contact-free Brillouin microscopy combined with the microfluidic vascular chip and the schematic of Brillouin measurements across the long axis of cells. **e**, Time-dependent increase in Brillouin shift in response to USS and OSS (n=5 for each condition). **f**, Schematic representation of cells subjected to USS with YAP showing initial nuclear localization followed by cytoplasmic retention as the flow duration increases, and cells subjected to OSS with continuous YAP nuclear localization. Quantification of YAP nuclear-cytoplasmic ratio (YR) in response to USS and OSS (3 independent experiments, n > 80 cells). **g**, Linear regression of YR with respect to total fluorescence intensity of actin cap under 24 hours USS and OSS (p<0.01). **h,i**, Comparison of LatB untreated, 30nM LatB treated, and 60nM treated cells under 24 hours USS, and the static control in total fluorescence intensity of actin cap and YR (3 independent experiments, n > 80 cells). **j**, Schematic representation of cells in the laser ablation experiment treated with 30nM, 60nM, and 120nM LatB. The recoil angle in response to laser ablation (n=5 for each condition) and the corresponding stress within the basal conventional fibers. All data are shown as mean□±□s.d. *p<0.05; **p<0.01; ***p<0.001; ****p<0.0001; NS, not significant.

The percentage of cells with actin cap (Extended Data Fig. 2a,b), the total fluorescence intensity of actin cap and the chromatin compaction increased with flow in a time-and magnitude-dependent manner under both 12 dyne/cm^2^ (Fig. 1a-c) and 4 dyne/cm^2^ unidirectional shear stress (Extended Data Fig. 2c-e). Considering the connection between nuclear stiffness and chromatin compaction^18^, we employed Brillouin microscopy for label-free, contact-free temporal measurements of nuclear stiffness (Fig. 1d), revealing increased stiffness under both unidirectional conditions (Fig. 1e, Extended Data Fig. 2f). Furthermore, YAP initially entered and subsequently gradually exported from the nucleus (Fig. 1f, Extended Data Fig. 2g), in line with previous studies^11^. In contrast, oscillatory shearing (1Hz, 0 ± 12 dyne/cm^2^) resulted in more gradual and less extensive actin cap formation (Fig. 1b), with slightly reduced chromatin compaction and nuclear stiffness (Fig. 1c,e). Notably, YAP remained localized in the nucleus (Fig. 1f), corroborating prior findings^13^.

**Fig. 2.**
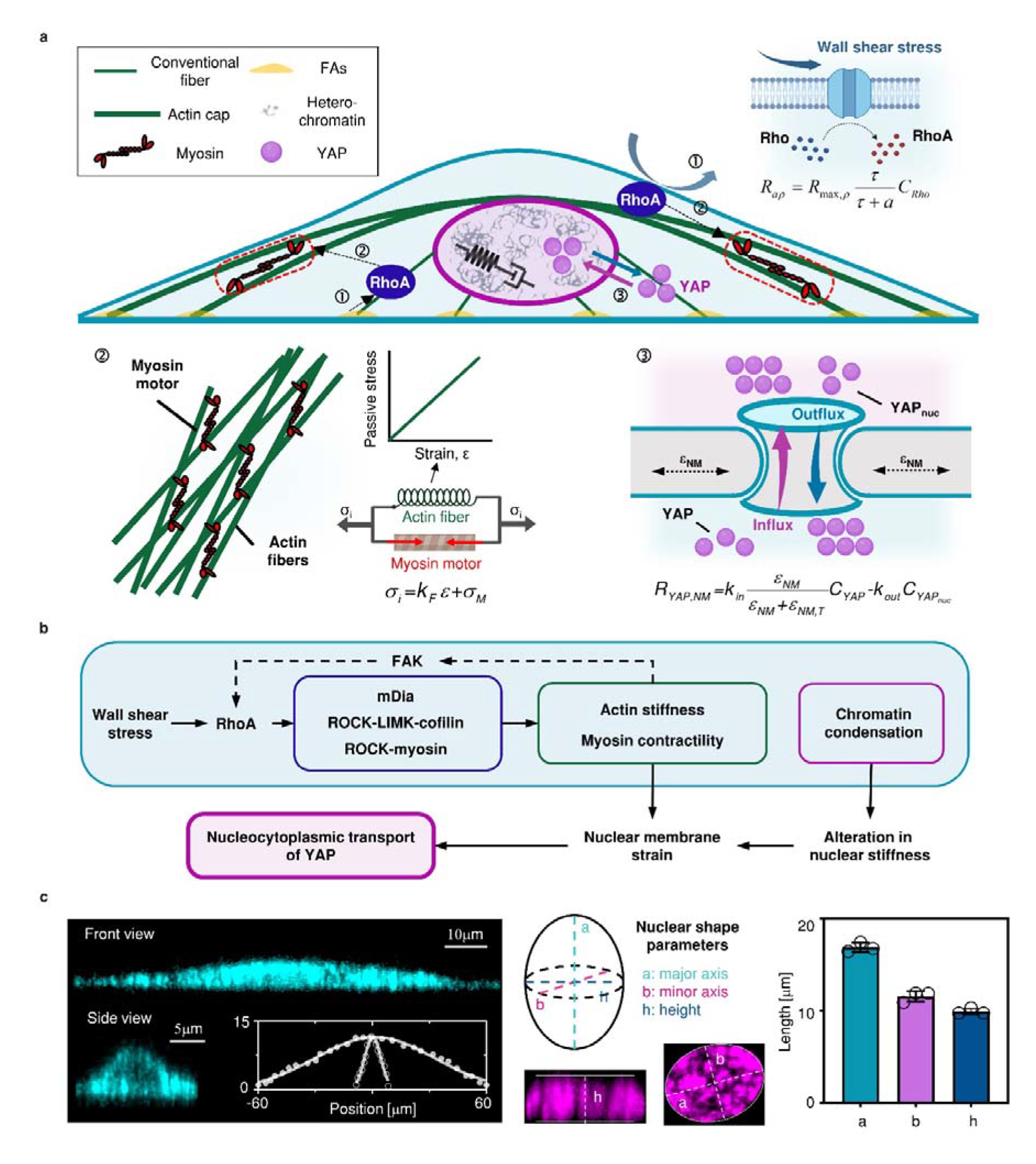
The schematic of three-dimensional mechanochemical model. **a**, The components considered in the model, including the cytoplasm, conventional fiber, actin cap, nucleus, and biochemical pathways centered on RhoA to regulate the transport of YAP (FAs indicates the focal adhesion in the figure). **b**, Detailed biochemical pathways involved in the model, including the flow-activation of RhoA, cytoskeletal dynamics regulation pathways downstream of RhoA involving ROCK, mDia, LIMK and Cofilin, and YAP transport. **c**, Geometric parameters obtained from cytoplasm (stained with CM-Dil), fitted by the Gaussian curve, and nuclear images (stained with DAPI), described by the length of major axis, minor axis, and height.

We furthermore conducted correlation matrix analysis among the mean YAP nuclear-cytoplasmic ratio (YR), actin cap, and nuclear stiffness for each flow condition. We identified a significant negative correlation between YR and nuclear stiffness (Extended Data Fig. 2h). Additionally, we observed a negative correlation between YR and the total fluorescence intensity of the actin cap (P=0.06). To better understand the role of the actin cap, we investigated the relationship between the total fluorescence intensity of the actin cap and YR for individual cells under each flow conditions. Our results demonstrated that cells with higher actin cap intensity displayed lower YR values under both unidirectional and oscillatory shear stress (Fig. 1g, Extended Data Fig. 2i).

To validate the role of actin cap, we suppressed its formation using low doses of latrunculin B (LatB, <60nM) under 12 dyne/cm^2^ unidirectional shear stress^19, 20^. Upon inhibiting (Fig. 1h, Extended Data Fig. 3a), a significant increase in YR was observed after 24 hours of shearing compared to the group without actin cap inhibition (Fig. 1i). Previous research indicates that low LatB concentrations preferentially inhibit actin cap than basal conventional actin fibers^20^. Our immunofluorescence staining also demonstrates that LatB concentrations below 60nM preserve conventional fiber structure (Extended Data Fig. 3c). In addition, we conducted laser ablation experiment and revealed minimal impacts on recoil angle of conventional fibers for 30nM and 60nM LatB treatments (Fig. 1j). Subsequently, we utilized the vertex model (see Methods) to simulate the laser ablation experiments and quantitatively corroborated the relatively minor effects on the contractile stress of conventional fibers at these concentrations (<9% for 30nM and <36% for 60nM, Fig. 1j). In contrast, 120 nM LatB treatment substantially reduce the contractile function of conventional fibers, guiding our focus to treatments below 60nM.

**Fig. 3.**
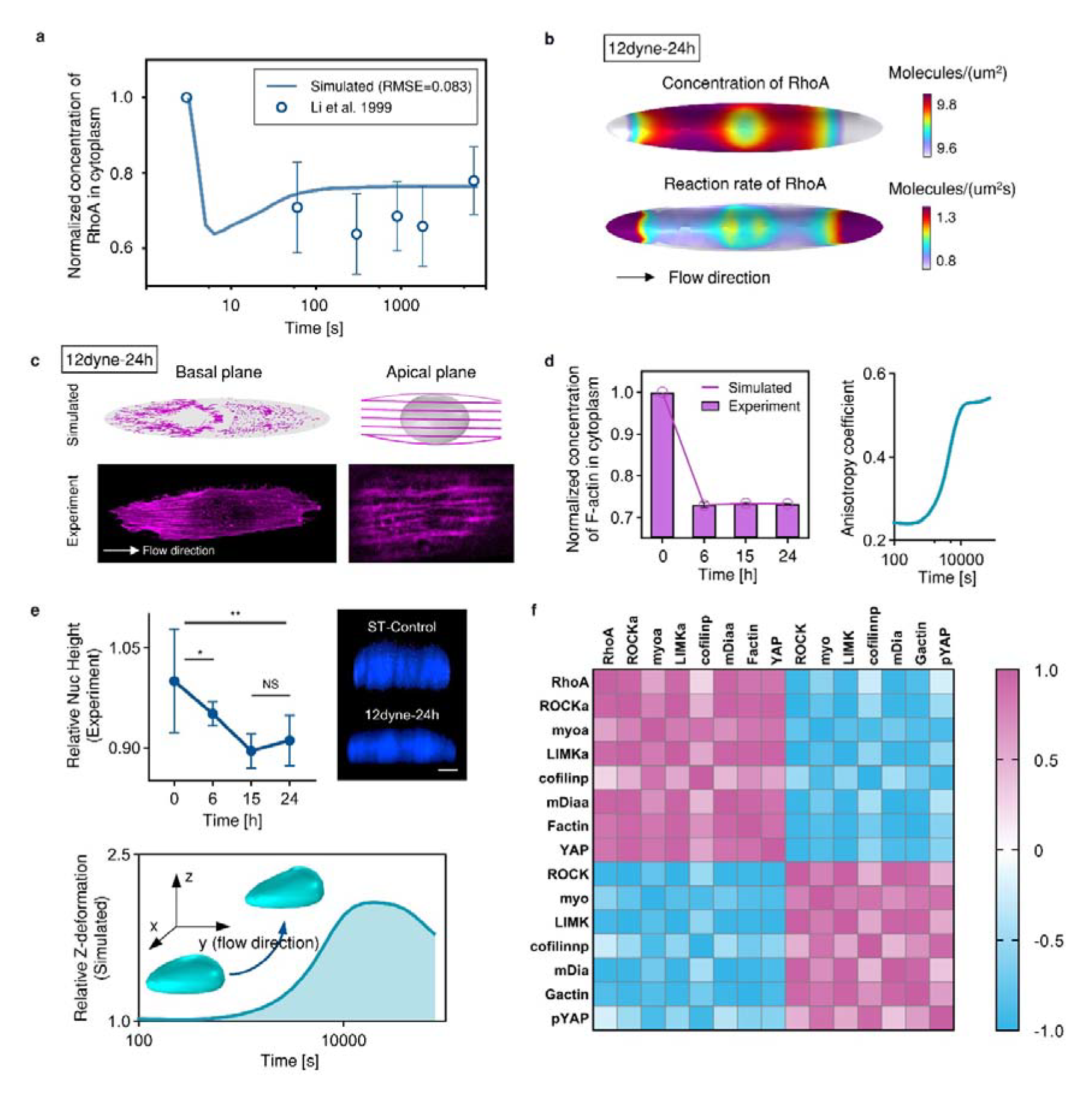
The prediction of mechanochemical behaviors involved in shear stress mechanotransduction. **a**, Comparison between the model simulated RhoA concentration in the cytoplasm and experimental measurement^21^. **b**, The spatial distribution of RhoA concentration and reaction rate at the cell membrane under USS. c, Experimental and simulated organization of actin fibers, including conventional fibers in the basal plane and actin cap in the apical plane. **d**, Experimental and simulated formation of conventional fibers, and the simulated polarization of conventional fibers described by the anisotropy coefficient calculated by the ratio of stress parallel to flow versus perpendicular to flow. **e**, Experimentally measured nuclear height changes under USS (3 independent experiments, n>80; *p> 0.05; *p<0.05; **p<0.01; NS, not significant. mean□±□s.d.). Simulated nucleus’s z-direction deformation under USS. **f**, Correlation matrix analysis of the concentrations of cascade signaling pathways involved in the model, including RhoA, and the activated (inactivated) ROCK, myosin, LIMK, cofilin, mDia, F-actin, and YAP.

## Mechanochemical model to analyze the flow shear stress transmitted to nucleus regulating nucleocytoplasmic transport of YAP

A three-dimensional cell is included in the model, which comprises the cytoplasm, conventional fibers, actin cap, and nucleus (Fig. 2a). The cell is embedded in a microfluidic channel (Extended Data Fig. 4a). Using immunofluorescence staining with CM-Dil, phalloidin, and DAPI, the model effectively captures these subcellular components (Fig. 2c). We considered both the involved biochemical pathways and mechanical transmission, including the activation of RhoA (numbered as ① in Fig. 2a), cytoskeleton reorganization (numbered as ② in Fig. 2a), and YAP nucleocytoplasmic transport (numbered as ③ in Fig. 2a) in this model.

**Fig. 4.**
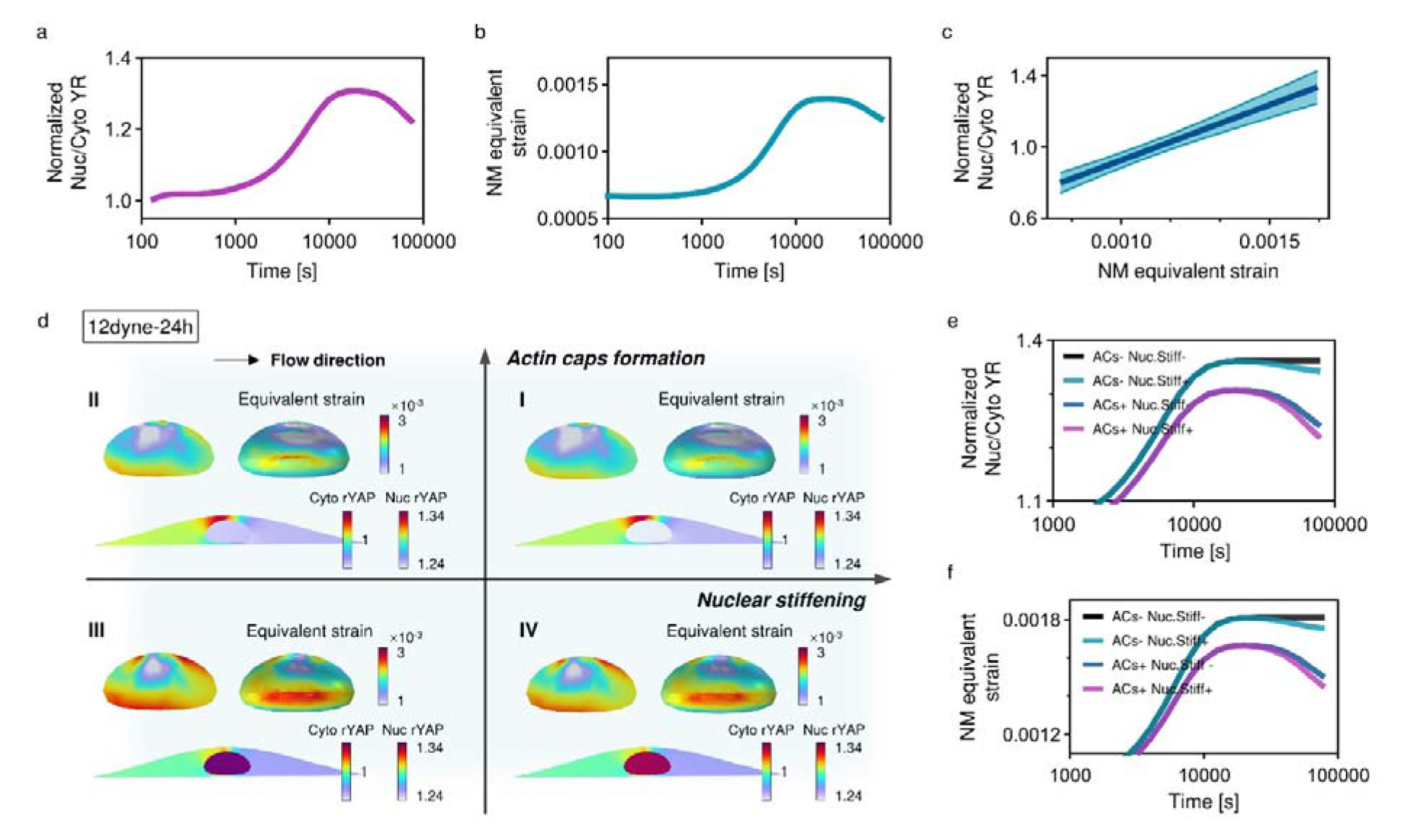
The roles of actin cap and nuclear mechanics in regulating YAP biphasic transport. **a,b**, Simulated YAP nuclear-cytoplasmic ratio (YR) and nuclear membrane (NM) equivalent strain under USS of 12 dyne/cm^2^ in 24 hours. **c**, Linear regression of YR with respect to NM equivalent strain (p<0.001). **d**, Individual effects of actin cap and nuclear stiffening in shear stress mechanotransduction and YAP nucleocytoplasmic shuttle (t=24 hours, Quadrant I: actin cap+, nuclear stiffening+; Quadrant II: actin cap+, nuclear stiffening-; Quadrant III: actin cap-, nuclear stiffening-; Quadrant IV: actin cap-, nuclear stiffening+). **e,f**, The explicit effects of actin cap and nuclear stiffening in surface-averaged YR and NM equivalent strain under USS, illustrating the dominating effects of actin cap and the collaborating effects of nuclear stiffening on the biphasic transport of YAP.

## i. Flow-induced RhoA activation

Firstly, flow shear stress activates integrins and G protein-coupled receptors at plasma membrane^21^, leading to RhoA activation, expressed by Hill equation:

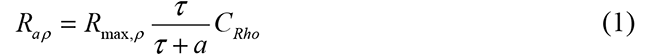

where *R_max,ρ_* represents the maximum activation rate, τ denotes shear stress magnitude, *C_Rho_* is Rho concentration, and *a* is shear stress value when the activation rate of RhoA reaches *R_max,ρ_*/2. Additionally, stress-transmitting actin fibers at the focal adhesion convert FAK to RhoA (Extended Data Fig. 5a)^22^. Our model’s simulated RhoA concentration shows good agreement with the experimental measurement^21^ (Fig. 3a).

**Fig. 5.**
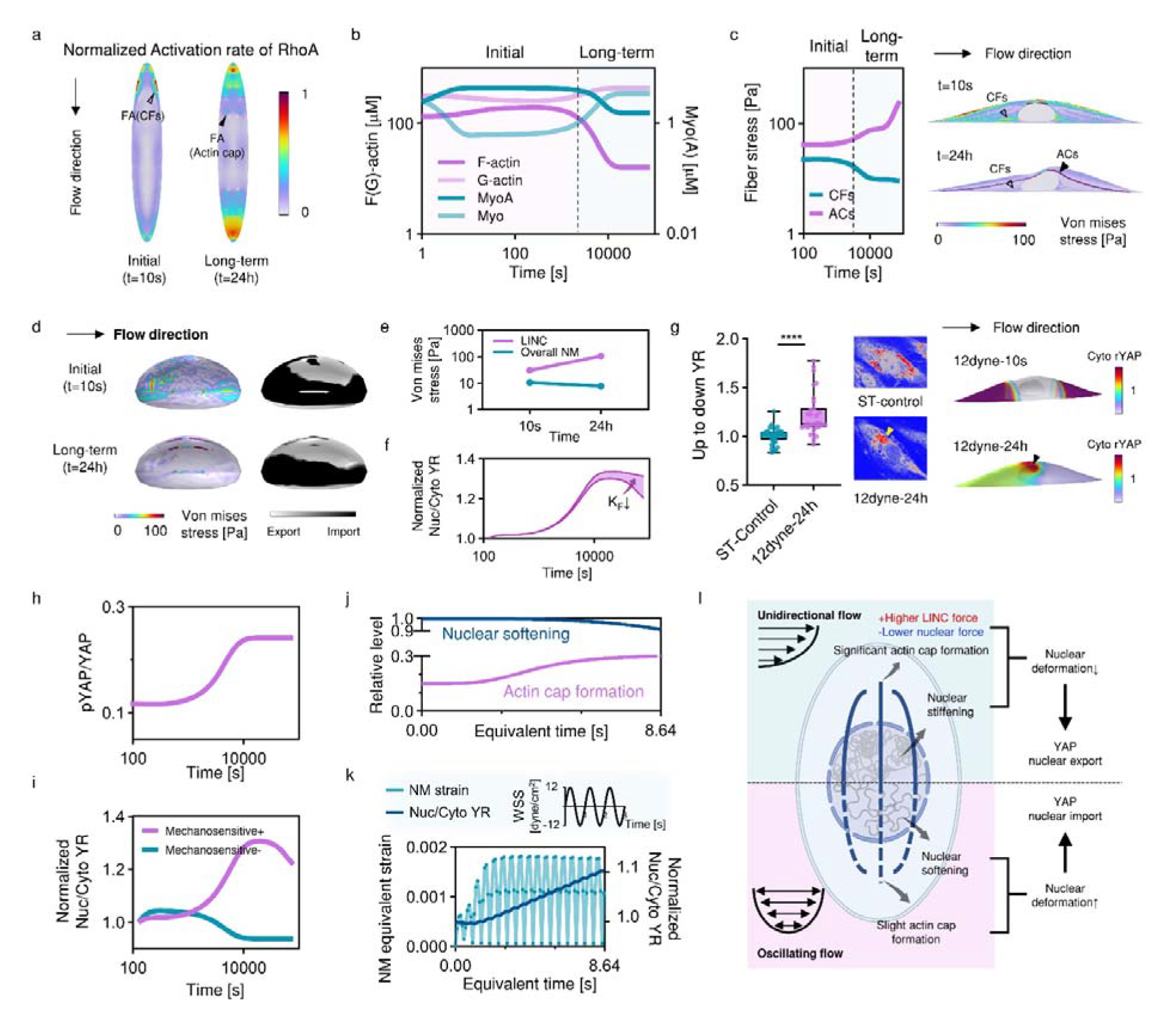
Mechanical mechanisms of flow shear stress-driven YAP spatiotemporal nucleocytoplasmic shuttling. **a**, Normalized RhoA activation rate at the basal plane at the initial (t=10s) and end state (t=24 hours) of USS (FA: focal adhesion, CFs: conventional fibers). **b**, Simulated spatial averaged concentration of F-actin, G-actin, myosin, and activated myosin during USS. **c**, Averaged Von Mises stress in conventional fibers and actin cap during USS, and the Von Mises stress distribution inside the cell at the initial (t=10s) and end state (t=24 hours) of USS. The deformation is visualized and expanded by a factor of 100. **d**, Nuclear stress distribution and YAP import/export flux at the initial (t=10s) and end state (t=24 hours) of USS. **e**, Nuclear stress in the actin cap-associated LINC and nuclear membrane (NM). The stress at actin cap-associated LINC increases as USS prolonged, while the stress at the overall NM significantly reduces. **f**, The effect of actin cap stiffness (KF) on the biphasic response of YAP transport while maintaining its contractile stress. **g**, Experimental YAP spatiotemporal distribution in ST control and USS of 12 dyne/cm2 for 24 hours (3 independent experiments, n > 30 cells. p<0.0001. mean□±□s.d.). Arrowhead indicates the unsymmetrically polarized spatial distribution of YAP in the cytoplasm. **h**, The activation of YAP, described by the ratio of pYAP to YAP, with respect to USS time. **i**, The comparison of artificially fully biochemical regulation of YAP and mechanochemical regulation of YAP. **j**, Relative changes in actin cap and nuclear stiffness under OSS. **k**, NM strain exhibits an oscillatory trend and YR increases as the duration of OSS prolongs. **l**, Prolonged USS leads to significant formation of actin cap and an increase in nuclear stiffness, while OSS results in slight actin cap formation accompanied by a decrease in nuclear stiffness, collaboratively regulating YAP transport.

## ii. Cytoskeleton reorganization

Following RhoA activation, rho-associated kinases (ROCK), formin mDia1 (mDia), LIM kinase (LIMK) and Cofilin pathways are activated (Fig. 2b). The transporting processes of these components are governed by reaction-diffusion equations (Supplementary Table 1), with coefficients obtained from recent biochemical models^22, 23^. These pathways control cytoskeletal stress, which depends on active and passive constituents, such that σ*_i_=K_F_*ε*+*σ*_M_*, where *K_F_* is effective passive cytoskeletal stiffness and σ*_M_* is the myosin-dependent contractile stress^16, 24^ (full derivation outlined in Supplementary Table 2). The formation of conventional fibers is modeled as a first-order function regard to F-actin concentration, while the formation of actin cap is directly derived from the experimental measurements. Our model demonstrates that the asymmetric spatial distribution of RhoA concentration and generation rate (Fig. 3b) corresponds to more pronounced polar remodeling of basal conventional fibers facing the flow (Fig. 3c). Furthermore, our model successfully simulates the time course of basal conventional fiber formation and polarization under flow shear stress (Fig. 3d, Extended Data Fig. 5b).

## iii. YAP nucleocytoplasmic transport

Lastly, we modeled a set of reactions accounting for the interplay between F-actin, myosin, and YAP dephosphorylation^12, 25^. The rate of YAP dephosphorylation are modeled as a product of F-actin and myosin concentration^22^, while YAP nuclear localization or cytoplasmic retention is determined by the export and import rates of dephosphorylated YAP. In this model, the YAP nuclear import rate is set as a Hill function of nuclear membrane equivalent strain, reflecting nuclear pore stretch, while the YAP export rate follows a first-order rate^6, 7, 22^. Our model aligns well with experimentally measured nuclear heights, demonstrating a decrease in nucleus height under flow shear stress, followed by a rebound after 15 hours (Fig. 3e).

Using the proposed mechano-chemical model, we reproduced correlations among the involved signaling molecules: positive correlations between RhoA and downstream activated ROCK and myosin, positive correlations between RhoA and downstream mDia and F-actin, and correlations between RhoA, activated ROCK, LIMK, cofilin and F-actin (Fig. 3f). Full model parameter definitions, values, and the sensitivity analysis of key parameters are provided in Supplementary Table 1-2, and Extended Data Fig. 5d.

## Combined effects of actin cap formation and nuclear mechanics on biphasic YAP nucleocytoplasmic shuttling under unidirectional flow shear stress

Initially, we utilized the Hill function to fit the relationship between flow time, shear stress magnitude, actin cap formation and Young’s modulus of the nucleus (Extended Data Fig. 4b-c). The formation of actin caps (ρ) was fitted from the actin cap present cells (Extended Data Fig. 2a) while the Young’s modulus of the nucleus was obtained from a log-log linear relationship^26^ based on experimentally measured Brillouin shifts (Fig. 1e). Our simulation results showed an increase in YR, followed by a gradual decline under unidirectional shear stress condition (Fig. 4a, Supplementary Video 1). Additionally, we observed faster YAP transport process under 12 dyne/cm² of unidirectional shear stress compared to 4 dyne/cm² (Extended Data Fig. 6a), corroborating experimental observation (Fig. 1f). Moreover, we identified a similar biphasic pattern for nuclear membrane strain (Fig. 4b) that positively correlated with YR (Fig. 4c), in line with previously reported experimental findings^6, 7^.

**Fig. 6.**
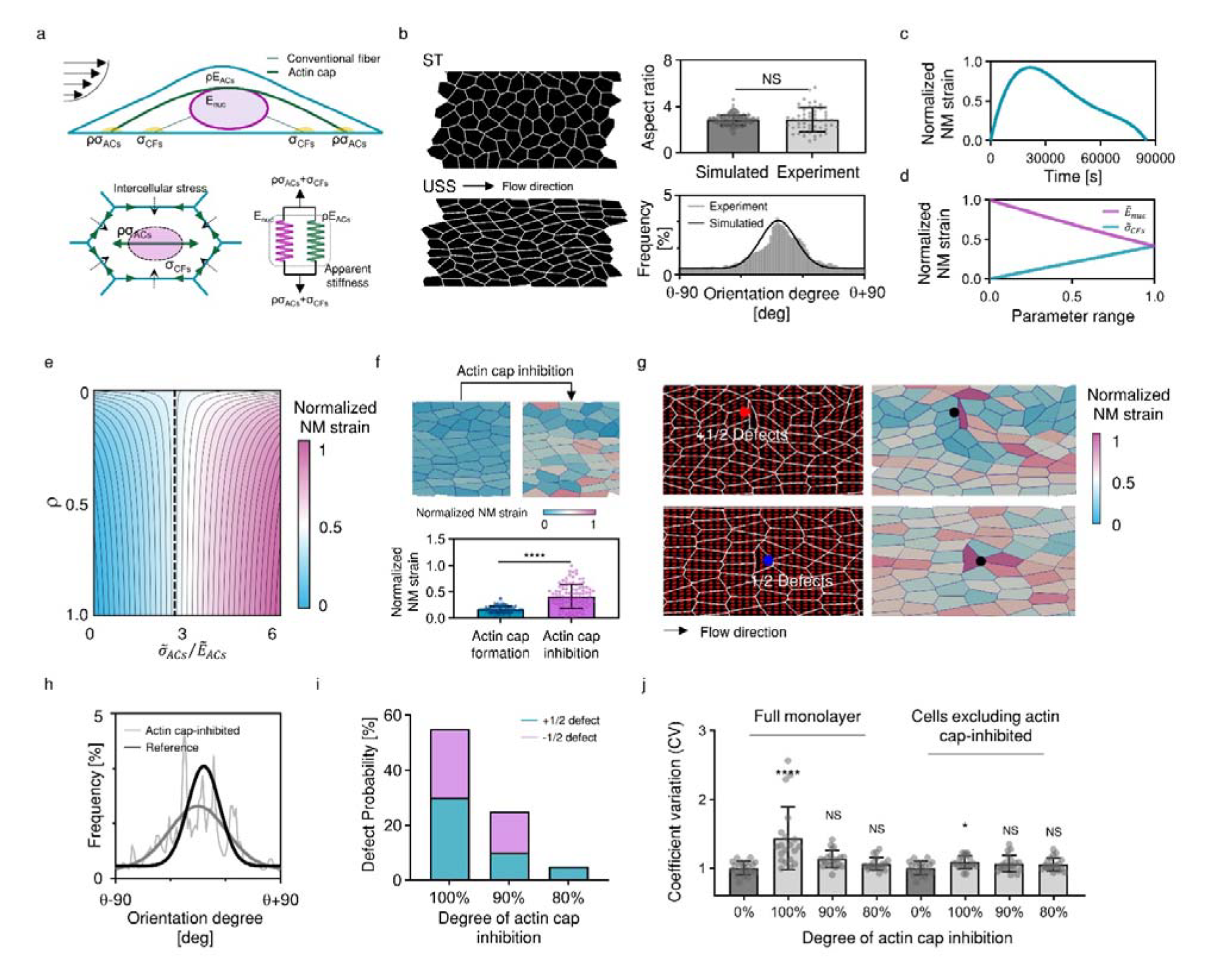
Regulatory role of actin cap in cell monolayer behavior and nuclear membrane strain. **a**, Schematic of the vertex model considering the formation of actin cap and the minimal model to assess the collective cells’ nuclear membrane strain. **b**, Static and USS-induced alignment, and elongation of endothelial cell monolayers simulated by the vertex model. The comparison of experimental and simulated cell aspect ratio and polarization (the flow direction is denoted as θ). **c**, Minimal model-calculated nuclear membrane strain over the duration of USS. **d**, Dimensionless nuclear stiffness 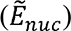 and conventional fiber stress 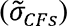 and their impact on nuclear membrane strain (nuclear stiffness and conventional fiber stress are dimensionless based on the three-dimensional modeling results, with 1.45 MPa and 560 Pa, respectively). **e**, The influence of actin cap on nuclear membrane strain depends on the ratio between its stiffness 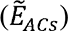 and stress 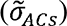. **f**, The effect of actin cap inhibition on collective cell nuclear membrane strain (n > 80, p < 0.0001, mean ± s.d.). **g**, The inhibition of actin cap in individual cells and its potential role in inducing topological defects and spatially heterogeneous distribution of nuclear membrane strain (red dots represent +1/2 defects, blue dots represent −1/2 defects). **h**, The effects of actin cap inhibition in individual cells on collective cell polarization. **i**, The varying degrees of actin cap inhibition in individual cells impact the formation and probability of different defect types (n = 20 in each condition). **j**, The varying degrees of actin cap inhibition in individual cells influence the spatially heterogeneous distribution of nuclear membrane strain, including and excluding the cells with inhibited actin cap (n = 20 in each condition, ****p < 0.0001; *p < 0.05; NS, not significant).

Next, we examined whether the coupling effects of actin cap and nuclear mechanics dominates YAP nucleocytoplasmic shuttling by selectively inhibiting these factors. Firstly, we compared the nuclear membrane strain and YR with various conditions of actin cap and nuclear mechanics inhibition at the same flow time of 24 hours. Our simulations predict that nuclear stiffening alone (actin cap-, nuclear stiffening+, Fig. 4d Quadrant IV) induces a higher nuclear membrane strain compared to the coupling effects (actin cap+, nuclear stiffening+, Fig. 4d Quadrant I), resulting in a stronger trend of YAP nuclear localization. On the other hand, the effect of actin cap alone (actin cap+, nuclear stiffening-, as shown in Fig. 4d Quadrant II) is consistent with the effect of nuclear stiffening alone but is not significant. Conversely, when both factors are inhibited (actin cap-, nuclear stiffening-, Fig. 4d Quadrant III), the nuclear membrane strain increases significantly, promoting higher YAP nuclear concentration.

To further understand the dynamics of YR and nuclear membrane strain, we calculated their spatial averaged values in 24 hours. Our results showed that inhibiting both actin cap and nuclear stiffening decreased YAP export and the nuclear membrane strain decreasing tendency under prolonged flow. Notably, when both factors are simultaneously suppressed, the decreasing trend of YR and nuclear membrane strain completely disappears (Fig. 4e,f). These findings suggest that the biphasic response of nuclear membrane strain and YAP to unidirectional shear stress is primarily governed by the coupling effects of actin cap and nuclear mechanics.

## Mechanism in spatiotemporal mechanotransduction of flow shear stress-driven YAP nucleocytoplasmic shuttling

Upon the onset of unidirectional shear stress, the actin cap only slightly forms. Instead, RhoA and the upstream stress sensitive FAK were mainly activated at focal adhesions linked to conventional fibers (Fig. 5a, Extended Data Fig. 5a). Multiple pathways, such as mDia, ROCK, LIMK, and Cofilin (Extended Data Fig. 5c), interacted to rapidly increase F-actin levels and myosin activation within conventional fibers. These levels stabilized during the initial stage of flow (Fig. 5b), resulting in highly contractile fibers capable of generating increased stress (Fig. 5c, hollow arrows). The subsequent rise in intracellular stress facilitated the dispersal stretching of the nuclear membrane, promoting YAP import into the nucleus (Fig. 5d).

As the flow duration progressed, the formation of actin caps became prominent. Concurrently, RhoA and FAK activation shifted from conventional fiber focal adhesions to the actin cap (Fig. 5a, Extended Data Fig. 5a), leading to reduced stress in conventional fibers (Fig. 5c, hollow arrows) and increased stress within the actin cap (Fig. 5c, solid arrows, Supplementary Video 2). The actin cap not only exerts contractile stress but also limits nuclear deformability. This resulted in concentrated high levels of stress on the LINC protein complex, while the overall nuclear membrane stress was reduced (Fig. 5d-e). The combination of decreased stress and flow-induced nuclear stiffening promotes YAP export from the nucleus (Fig. 5d). When we artificially decrease the stiffness of actin cap while maintain its contractile stress, the biphasic response of nuclear membrane strain and YAP export are inhibited (Fig. 5f). Furthermore, force applied by the actin cap on the nuclear surface leads to a higher propensity for YAP nuclear import at this location, causing cytoplasmic YAP accumulation (Fig. 5g). In addition, the asymmetric polarity of flow results in a higher YAP concentration in the cytoplasm facing the flow, consistent with our experimental observations (Fig. 5g).

In addition to mechanical regulation, the chemical activity of YAP has been demonstrated to contribute to its transport process^5, 13^. Our model results indicate that the ratio of phosphorylated YAP (pYAP) to dephosphorylated YAP increases as unidirectional shear stress persists (Fig. 5h), consistent with previous experimental studies^5, 13^. To elucidate the role of decreased YAP activity in YAP export under long-term flow, we suppressed the mechanosensitive aspect of YAP transport in our model. Interestingly, our simulations showed a rapid YAP nuclear export reaching a stable value (Fig. 5i), differing from the experimental results on YR dynamics (Fig. 1f), suggesting mechanical regulation is a crucial mechanism of YAP transport.

We further utilized our model to investigate the impact of oscillating shear stress on nuclear membrane deformation and its effect on YAP transport. We simulated the process of actin cap formation and nuclear softening over a 24-hour period (86,400s) condensed into an 8.64-seconds simulation (Extended Data Fig. 4b,c, Fig. 5j). Our results demonstrate that, due to the highly contractile nature of conventional fibers and the relatively low actin cap formation and nuclear softening compared to unidirectional shear stress, cells experience a significant increase in nuclear membrane strain with each oscillating period. This substantial nuclear membrane deformation consistently promotes YAP nuclear import (Fig. 5k). Furthermore, artificially enhancing actin cap formation and inhibiting nuclear softening during oscillating shear stress led to a reduction in both nuclear membrane strain and YAP nuclear entry over time (Extended Data Fig. 6b).

## Regulation of cell monolayer behavior by actin cap formation

In cell monolayers, topological defects of cell alignments have been reported to influence spatial varying YAP activity, mediating cell death and extrusion processes^27, 28^. Considering the close relationship between actin stress fiber and cell alignments under flow, and the impact of actin reorganization on collective cellular mechanotransduction^29, 30^, we integrated our single-cell mechanochemical model with the vertex model to investigate the regulatory role of actin cap in cell monolayer behavior with focus on nucleus membrane strain, which is the key factor for YAP dynamics. To achieve this, we modified the vertex model to consider the flow-induced formation of actin cap. The cell shapes and stress in conventional fibers (σ_CFs_) are determined by their packing topology^31^. The stress in actin cap (σ_ACs_) is implemented by an anisotropic bulk stress within each cell (Fig. 6a, full derivation outlined in Methods). We found that upon actin cap forming, the vertex model successfully reproduced the experimentally measured endothelial cell aspect ratio and alignments under flow (Fig. 6b). To achieve a relatively general and concise model of various mechanical stimulations such as flow shear, substrate stretching and stiffness, we derived a minimal model of nuclear membrane strain based on our 3D modeling and integrated it with the vertex model (Fig. 6a), which is expressed as:

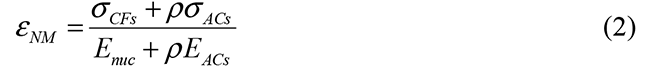

Here, *E_nuc_* represents nuclear stiffness, *ρ* denotes actin cap formation, and *E_ACs_* corresponds to actin cap stiffness. The apparent stiffness, *E_nuc_+*ρ*E_ACs_*, is determined by both the nucleus and the actin cap. To validate the reliability of equation (2), we applied the 3D model’s simulated time sequences of stress and successfully reproduced the initial rise and decline in nuclear strain (Fig. 6c). In addition, Equation (2) captures the relationship between nuclear stiffness, conventional fibers stress, and nuclear membrane strain (Fig. 6d), consistent with our results and previous findings^6, 7, 22, 23^. The influence of actin cap on nuclear membrane strain depends on the ratio between its stiffness and stress (Fig. 6e). A larger σ*_ACs_* to *E_ACs_*ratio leads to increased nuclear strain as actin cap form, potentially corresponding to actin cap formation under short-term mechanical stimuli, as observed in periodontal ligament cells responding to flow within 1 hour^25^ and cells exposed to substrate stretching and stiffness^32^, leading to YAP nuclear import. In contrast, a smaller ratio decreases nuclear strain, potentially reflecting long-term actin cap formation, such as in endothelial cells responding to unidirectional flow and cells experiencing prolonged stretching, leading to YAP nuclear export^33^.

The vertex model results show that the inhibition of actin cap in cell monolayer results in an increased nuclear membrane strain (Fig. 6f), which agrees with our experimental results of actin cap inhibition promotes YAP nuclear import (Fig. 1i). Then, we selectively inhibited actin cap formation in individual cells and found that the localized actin cap irregularities affect collective cells alignments (Fig. 6h). Notably, the simulation results predicted that the actin cap irregularities can induce topological defects and spatially heterogeneous distribution of nucleus membrane strain (Fig. 6g, Extended Data Fig. 6c). The defects probability and coefficient of variation for spatial distribution of nucleus membrane strain exhibited a significant increase when the target cell underwent complete actin cap inhibition. This increase progressively diminished as the degree of actin cap inhibition decreased (Fig. 6i,j). Collectively, our vertex model analysis predicts the pivotal role of actin cap in modulating collective cell alignments and spatial distribution of nucleus membrane strain within cellular monolayers.

Taken together, our study demonstrates how the nucleocytoplasmic shuttling of YAP under flow shear stress is controlled by the coupling between the actin cap and nuclear mechanics. Through the combination of microfluidics and Brillouin microscopy, we identified a negative correlation between YAP nuclear-cytoplasmic ratio (YR), actin cap, and nuclear stiffness under various flow types. Given the multifaceted effects of LatB treatment, such as calcium activity^34^ and chromatin compaction alteration^19^ (Extended Data Fig. 2b), we developed a mechanochemical model to investigate the explicit role of actin cap and nuclear mechanics. We demonstrated that actin cap concentrates stress within the LINC and provides stiffness constraint to reduce the nuclear deformation, echoing recent findings in substrate stretching fibroblast cells^2^. Furthermore, we found that nuclear stiffening decreases the deformation, regulating YAP nuclear export and aligning with recent studies on curvature-driven chromatin compaction and YAP localization^35^.

Our collective-cell model uncovers the significance of actin cap in regulating cell alignments, highlighting the potential induction of topological defects and heterogenous YAP distribution upon localized actin cap inhibition. Research has indicated that cytoskeletal disruption can result in cell extrusion or death^36^. Our results demonstrate that cells with inhibited actin cap exhibit increased nuclear stress (Fig. 6g, Extended Data Fig. 6c). This localized stress concentration raises the likelihood of cell extrusion or death^27, 37^. In addition, our analysis of nuclear strain suggests that cells with −1/2 topological defects may exhibit higher YR compared to cells with +1/2 defects (Extended Data Fig. 6c), consistent with the experimentally observed defect-induced YAP heterogenous activity^27^.

The interaction between actin cap and nucleus may represent a retention mechanism modulating gene expression. While our results focus on YAP, similar regulatory patterns may apply to other transcription factors, such as MRTF-A, β- catenin, and MyoD^6, 7^. Moreover, our model can be extended to study the mechanotransduction process involving RhoA-centered pathways under substrate stiffness or stretching^22, 23^. This offers a unified mechanical framework for understanding nucleocytoplasmic transport of transcription factors, with potential implications for multiscale mechanotransduction scenarios.

## Supporting information

Supplemental infomation

## Acknowledgments

This work was supported by the National Natural Science Research Foundation of China (grant no. 11827803, 31971244, 31570947, 32071311, U20A20390, and T2288101), the Fundamental Research Funds for the General Universities (KG16186101) and the 111 Project (B13003). Fig. 2a, and Fig. 5i are created with BioRender.com.

## Author contributions

T.M., X.L., and H.S. conceived, designed, and led the interpretation of the project. T.M., X.L., and Y.H. led the development of the overall model and theory. T.M., F.W., and S.L carried out the bio-optical measurements of cells. H.S., C.G., Z.L., D.Z., X.Z., K.L., K.H., J.Z., and L.W. carried out the biological and microfluidic experiments. M.W. provides the biological sample of cells. Y.H., W.H., X.C., and X.D. provided guidance on analyzing and interpreting the resultant of bio-optical measurements. T.M., X.L., P.W., and Y.F. designed the experiments. All authors contributed to the writing of the manuscript.

## Methods

### Cell culture

Human umbilical vein endothelial cells (HUVECs) were isolated from newborn umbilical cords, as detailly described in our previous study^38^. Informed consent was obtained for each donor and all processes were approved by the Beihang University Ethical Committee. The primary HUVECs were cultured in endothelial cell medium (ECM, ScienCell) with 5% FBS, 1% endothelial cell growth supplement (ECGS), and 1% penicillin/streptomycin solution, and incubated at 37 °C under 5% CO_2_. The cells were fed one time a week with a complete change of fresh culture medium and used between 2 to 6 passages.

### Microfluidic Vascular Chip Fabrication

The microfluidic device was fabricated out of polydimethylsiloxane (PDMS, Sylgard 184, Dow Corning) using standard soft lithography (Extended Data Fig. 1a,b). The microfluidic chip is made up of three layers containing PDMS channel layer, thin PDMS membrane middle layer and glass coverslip supporting layer. To make the PDMS channel layer, PDMS prepolymer with a 10:1 (w/w) mixture of PDMS base and curing agent was casted against a photolithographically prepared master with a designed microchannel made of photoresist (SU8-2075, MicroChem) on a silicon wafer, and then cured for 2 hours in an 80 [dry oven. The middle layer of thin PDMS membrane was produced by spin-coating PDMS prepolymer on silanized silicon wafers at 3500 rpm for 50s on a spin coater and baking on a hot plate at 180 °C for 30 minutes. The channel layer and thin PDMS membrane were cleaned with dusting compressed air tank, and then treated with oxygen plasma for 50 s to form covalent bonds between them. They were then placed in the oven for 10 minutes to strengthen the bonding. Finally, the device stripped from silicon was bonded to the glass coverslip in the same manner of oxygen plasma treatment.

Before cell seeding, microfluidic chips were sterilized for 1 hour by ultraviolet irradiation in the clean bench, and then the culture channels were coated with 125 μg/ml fibronectin. Endothelial cells were seeded into the channel and attached to the membrane surface for 1 hour at 37°C under static condition. In all experiments, the density of endothelial cells was controlled at ∼500 cells/mm^2^ to avoid the effects of cell density changes on YAP localization. The attached cells were then filled with ECM by two 3/32” barbed female Luer adapters (Cole-Parmer) inserted into the medium injection ports to provide nutrients, which the medium in the Luer adapter was changed every 12 hours.

### Control of Flow Shear Stress in Microfluidic System

The control of fluid flow system in the present study consisted of one micro-syringe pump (Lead Fluid), one electromagnetic pinch valve and a reservoir, which was used in our recent study^38^. Medium flow was provided synchronously by the programmable micro-syringe pumps. The electromagnetic pinch valve was used to switch the fluid flow between injecting into the microfluidic chip and withdrawing from the reservoir. To introduce flow into the microfluidic device, a sterile polypropylene barbed elbow fitting (1/16″, Cole-Parmer) connected to Teflon tubing was inserted into the inlet port of cell channel. The fluid flow was introduced from the chip’s outlet port into the reservoir and later withdrawn to the micro-syringe pump to create a highly efficient circulation system.

For the unidirectional flow, stable waveforms are used in the control system. For oscillatory flow, the microfluidic chips were connected to the circulation control system input a sinusoidal waveform often used for oscillating flow. The waveform of flow rate was also measured by flow sensor (ElveFlow, Extended Data Fig.1c).

### Immunofluorescence and Confocal Microscopy

Cells were fixed in 4% paraformaldehyde (PFA) for 10 minutes, permeabilized with 0.1% Triton X-100 in PBS at room temperature and blocked in 5% bovine serum albumin (BSA) for 1h. Cells were then treated by primary antibodies at 4°C overnight and incubated with appropriate secondary antibodies for 2 hours at room temperature. Specifically, YAP was stained mouse polyclonal anti-human YAP (sc-101199, 1:200, Santacruz) and Alexa Fluor 488 goat anti-mouse IgG H&L (ab150077, 1:200; Abcam). Phalloidin labeled with Alexa Fluor 488-conjugated phalloidin (YP0052L, 1:200; EVERBRIGHT) was used together with secondary antibodies. Nucleus were stained with 4′,6-diamidino-2-phenylindole dihydrochloride (DAPI; 1:1000; Sigma-Aldrich) for 10 min. For CM-DiI staining, the CM-DiI storage solution (C4060S,1:1000, Amresco) was added to the microfluidic device, and incubated for 5 minutes at 37[in a carbon dioxide incubator. The cells in the microfluidic device were then incubated in a refrigerator at 4□ for 15 minutes and washed with PBS solution for three times. All fluorescence images were collected by a laser scanning confocal microscopy (SP8X; Leica) with Leica Application Suite software (LAS X version 2.0.0.14332), using objective lens (NA = 0.6, magnification: 40×). When measuring nuclear heights, a 60× objective lens (NA = 1.4) was used to ensure a z-axis resolution of less than 200 nm.

### Latrunculin B Treatment

We achieved actin caps inhibition in endothelial cells by submerging cells in 30nM to 240nM Latrunculin B (LatB, Abcam) dissolved in DMSO solution (Sigma-Aldrich). To minimize the effects of LatB on conventional fibers rather than the actin cap, the time of LatB treatment was controlled under 1 hour. In flow experiments, LatB is added to the ECM medium at the corresponding concentration.

### Brillouin Microscopy Setup

A custom Brillouin microscopy was built to obtain biomechanical images of endothelial cells cultivated in the microfluidic vascular chip based on the recent protocol^39^. A 10mW single longitudinal mode CW 532nm laser (MSL-FN-532, CNI) was utilized as the light source to probe Brillouin scattering. The laser beam was focused into the center of the microfluidic channel by an objective lens (OBJ, NA = 0.6, magnification: 40×), providing a spot size of ∼0.5μm (transverse) and ∼2μm (axial). The backward scattered light was collected by the same OBJ, reflected at the polarized beam splitter (PBS), and coupled to the single-mode fiber (SMF). The microfluidic vascular chip was equipped on a 2D translational stage with 0.5μm resolution (SC-200, KOHZU Precision) to perform fast two-dimensional scanning.

The customized Brillouin spectrometer (Fig. 1d) has two stages with orthogonally oriented VIPA (Free Spectral Range 30GHz, LightMachinery). The light coming out of the fiber is coupled in the vertical direction to VIPA 1 through cylindrical lens CL1, and the spectral pattern output from VIPA 1 is projected onto mask 1 (M1) through cylindrical lens CL2. Then, spherical lens L1 couples the pattern on M1 to VIPA 2 in the horizontal direction, and spherical lens L2 projects the spectral pattern at the output of VIPA 2 onto mask 2 (M2). The spectral pattern is imaged by the CMOS camera (iXon, Andor) after relating through a 4f system consisting of spherical lenses and spatial filter, the spectral pattern is imaged by the CMOS camera (ORCA-Fusion, Hamamatsu). A Matlab software was developed to drive both the stage and the camera simultaneously and output the Brillouin shift data by automatically least square Lorentzian fitting the Brillouin spectra. In each cell nucleus region, we uniformly selected 10 positions for measurement and calculated the average value to obtain the overall nuclear Brillouin shift (ν_B_). To obtain the Young’s modulus of the nucleus used in the computational model, we first calculated the longitudinal modulus M’ according to^39^:

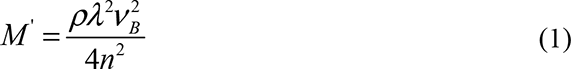

where ρ refers to the density of the nucleus (ρ=1350kg/m^3^), and λ is the wavelength of the incident light, n is the refractive index of the nucleus (*n*=1.38). Then, we transformed the longitudinal modulus M’ to the Young’s modulus E based on the log-log liner relationship^26^:

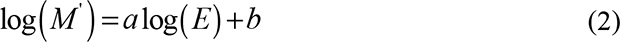

where *a* and *b* are the constant coefficients and equals to 0.081 and 9.37 according to the published study^26^.

### Preparation and mechanical measurements of gelatin methacryloyl (GelMA) Hydrogels

At 80 °C, different amounts of freeze dried GelMA macromer were dissolved in DPBS containing 0.5% (w/v) 2-Hydroxy-4’-(2-hydroxyethoxy)-2- methylpropiophenone (Sigma-Aldrich) as photoinitiator, yielding final GelMA concentrations of 5%, 7.5%, 15%, and 20% (w/v). After pipetting the prepolymer solution into a glass mold, it was subjected to 1000 mw/cm^2^ UV radiation (360-480 nm) for 180 seconds. In the uniaxial compression test, samples were made as cylinder specimens with diameter of 4 mm and thickness of 8 mm. Brillouin microscopy was then used to assess the stiffness of the samples, which was compared to the Young’s modulus derived from compression tests (Extended Data Fig. 1d,e).

### Laser ablation experiment

This experiment was performed using a femtosecond (fs) pulsed laser ablation system (852nm, 80 MHz repetition rate, InSight DeepSee). The basal layer of conventional fibers was ablated with 5 repeated cuts at 60 mW laser power. After the ablation, cells were allowed to relax for 30 minutes before being fixed. Immunofluorescence staining was then used to measure the recoil angle, calculated as the ratio of the short axis to the long axis of the opening ellipse, under different doses of LatB treatments.

### Measurements of Nuclear/Cytoplasmic Ratio of YAP

YR values were measured using image segmentation and quantifying the ratio of YAP intensity inside the nucleus to YAP intensity in cytoplasm. After projecting the maximum intensity of the image series onto a layer in ImageJ 1.5.3q^40^, we segmented the cytoplasmic YAP from the background using Cellpose, a pretrained deep learning-based segmentation method^41^. Then, we labeled the cell nucleus stained with DAPI via StarDist convolutional neural network^42, 43^. Then, the mean fluorescence intensity of YAP staining in the nucleus and cytoplasm in individual cells were measured respectively. Moreover, during the statistical analysis, we normalized the YR values relative to their respective static control groups to eliminate interference from systematic errors.

### Measurements of Chromatin Compaction with Nuclear DAPI Staining

Here, we used the microscopy-based approaches^44^ to assess the degree of heterogeneity of DNA signal across the nucleus of ECs exposed to shear stress by quantifing the coefficient of variation of DAPI intensity. After segmenting the nucleus with StarDist convolutional neural network^42, 43^, the coefficient of variation (CV) of individual nucleus is calculated as the ratio of standard deviation of the DAPI intensity values to the mean value of intensity of the nucleus. The segmentation of nucleus with StarDist was achieved in ImageJ 1.5.3q, and the standard deviation and the mean value of the DAPI intensity were calculated in MATLAB R2021a.

### Mechanochemical modeling of YAP spatiotemporal transport under flow shear stress

To investigate the mechano-chemical interaction among subcellular components under shear stress stimuli, a 3D finite element model (FEM) was constructed that connected fluid flow in the microfluidic channel, mechanical deformation, and biochemical cascade signaling transfer of adhering single cells. In this study, numerical simulations were carried out using the finite-element package COMSOL (Burlington, MA), which provides multi-physics coupling of the laminar flow, the solid mechanics, and the diluted species transport components (see simulation procedures in the Supplementary Notes). The model’s extracellular fluid properties were set to match our experimental setup in the microfluidic device. The cellular model was composed of three deformable components: cytoplasm, nucleus, and actin fibers (both conventional fibers and actin caps), the mechanical properties of which had previously been documented. The biochemical cascade signaling pathways were adapted from the previous numerical models of basal stiffness-mediated YAP transport^16, 22^, whose details can be found in the Supplementary Table 1,2.

### Vertex-based model simulating cell collective behavior

We employed the vertex model to represent the endothelial cell monolayer as a polygonal network, where interconnected polygons represent individual cells in contact^45^. The vertex dynamics follow a mechanical force balance, with each vertex position satisfying the overdamped dynamics equation:

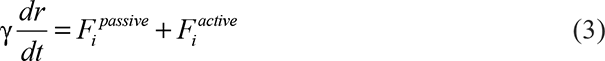

where *γ* denotes the friction coefficient between cells and the substrate. The passive force is described as 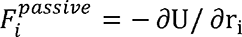, and the energy function *U* is defined as:

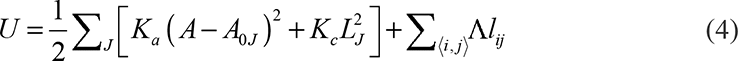

where *A*, *L_J_*, and *l_ij_* represent cell area, perimeter, and bond length, respectively. *r_i_* represents the position of vertex i. In addition, A_0J_ refers to the preferred cell area, *K_a_* and *K_c_* refer to the area and perimeter stiffness, and Λ represents the bond tension. Consequently, the three terms on the right-hand side correspond to area elasticity stress, perimeter elasticity stress, and intercellular tension, respectively. The active force 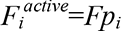 accounts for the traction forces of magnitude *F* exerted by the cells abutting at vertex i and in the average direction of their polarities *p_i_*=∑_<*α*|*i*>_ *p_α_*/*M_α_*. The cell polarity *p_α_* directs the traction force applied to cell α by the surrounding matrix. The time evolution of each cell polarity obeys the random rotation dynamics *dp_α_*/*dt*=□*(t)*^46^, with □*(t)* representing the white Gaussian noise.

For laser ablation experiment simulation, we eliminated four central cells in the middle to simulate the laser cutting. For flow shear stress simulation, we added a bulk stress in the vertex model to reflect the effect of shear stress on cell morphology based on the framework of Tlili^47^, which is assessed through the work of Lin et al.^48^. Aligning with the flow direction, the dimensionless bulk stress 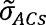 is transformed into the force applied to each vertex as:

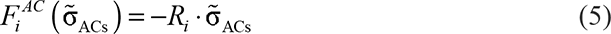

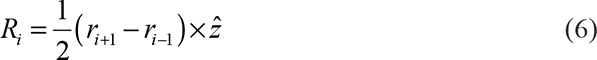

where *Ri* is the normal vector of the line *l_(i+1,i-1)_* connecting vertex *i+1* and *i-1*. *r_i+1_* and *r_i-1_* represent the position of vertex *i+1* and *i-1*, and 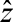 represents the unit vector,perpendicular to cell monolayer. The topological defects in the orientation field are detected by following the presented method^49^. In addition, the nuclear strain is calculated based on our proposed minimal model using the stress within actin cap and conventional fiber. The stress within conventional fiber is assumed to be the summation of cellular elasticity stress and intercellular tension. The computational method for the vertex-based model was implemented in CHASTE (University of Oxford, Oxford, UK)^50^. We used the following dimensionless values in the simulation to match the behavior of endothelial cell alignment: *A_0J_*=1, *K_a_*=1, *K_c_*=0.2, *P_0_*=- *Λ*/*K_c_*=3.6, 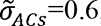, *F*=0.05, *γ*=1.

### Statistical Analysis

Data was analyzed using GraphPad Prism 9.0 software and were presented as mean ± SD. Sample numbers and experimental repeats were described in the figure legends. Statistical significance for samples was determined using two-tailed unpaired Student’s *t* test. Data were considered statistically significant if P < 0.05.

## Extended Data Figure

**Extended Data Fig.1.**
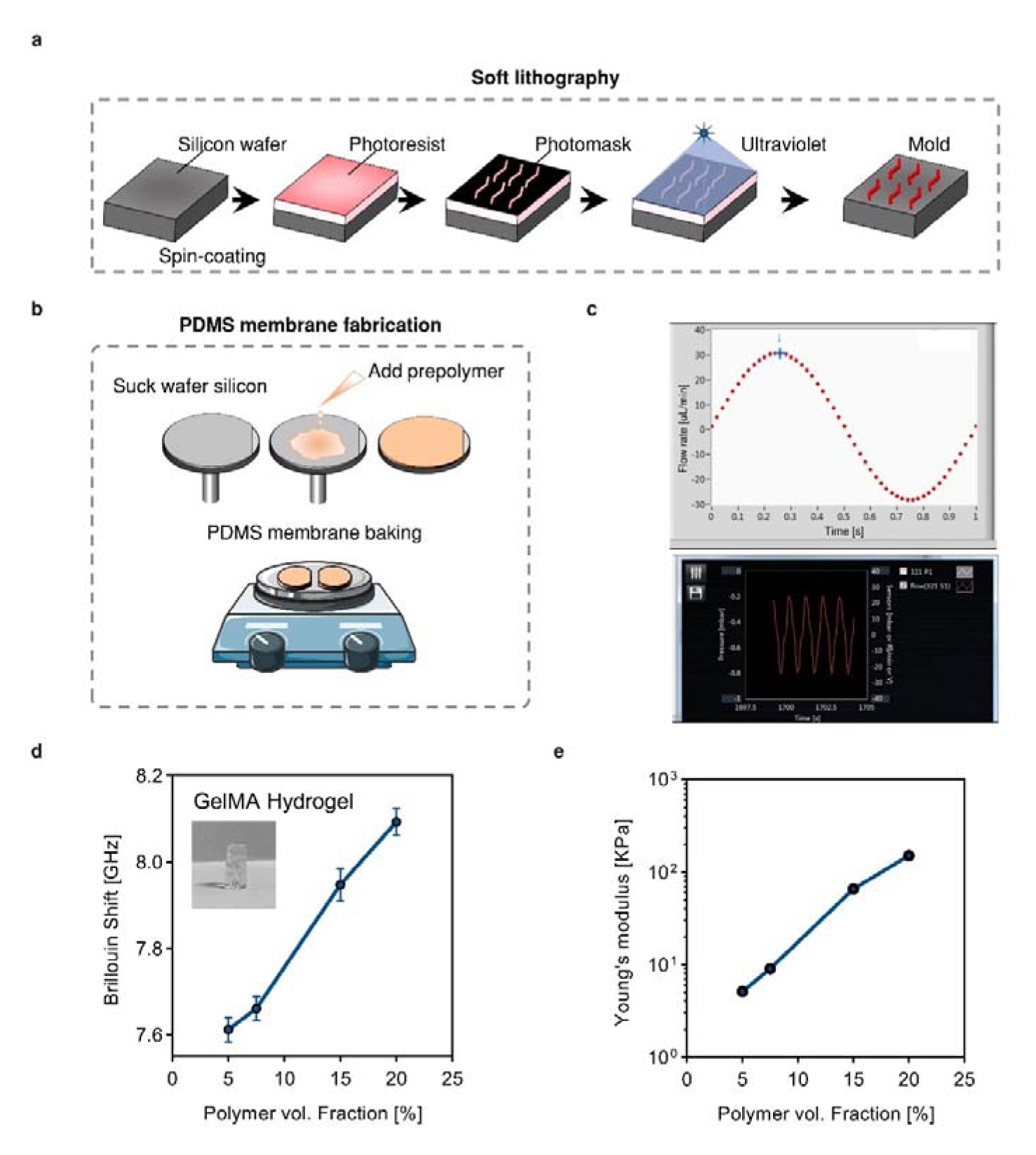
Microfluidic chip fabrication, flow control, and verification of Brillouin microscopy. **a,b**, Schematic diagram of soft lithography and PDMS membrane fabrication process. **c**, Representative illustration of flow control system in an oscillatory flow condition. **d,e**,The comparation of Brillouin shift measured by the Brillouin microscopy and the Young’s modulus measured by uniaxial compression tests. Both show the positive relationship with the GelMA fraction.

**Extended Data Fig.2.**
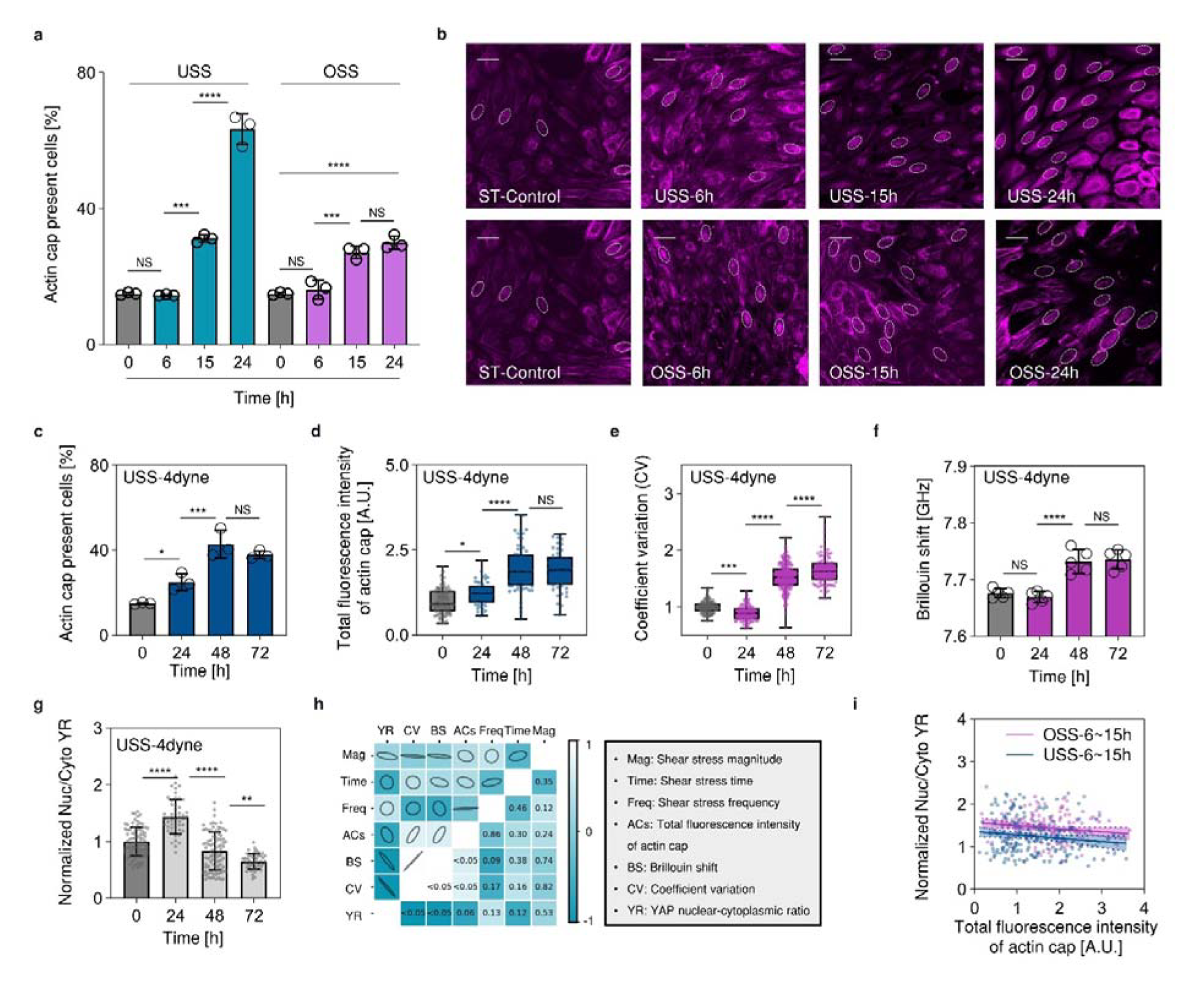
The dynamics of actin cap formation, nuclear stiffness, and YAP localization under flow shear stress. **a**, Flow-dependent formation of actin cap under USS of 12 dyne/cm^2^ and OSS of 0±12 dyne/cm^2^, 1Hz (3 independent experiments n > 80 cells). **b**, Apical view of actin cap formation under USS of 12 dyne/cm^2^ and OSS of 0±12 dyne/cm^2^, 1Hz. The actin cap present cells are marked with dashed lines (Scale bars, 40μm). **c-g**, Time-dependent variations in actin cap present cells, total fluorescence intensity, coefficient of variation, Brillouin shift, and YR under USS of 4 dyne/cm^2^ (3 independent experiments n > 80 cells, 5 independent experiments in Brillouin test). **h**, Correlation matrix analysis among shear stress magnitude, time, frequency, total fluorescence intensity, coefficient variation, and Brillouin shift. The upper half of the matrix displays correlation, while the lower half displays significance. **i**, Linear regression of YR with respect to total fluorescence intensity of actin cap under 6 to 15 hours USS of 12 dyne/cm^2^ and OSS of 0±12 dyne/cm^2^, 1Hz (p<0.01). All data are shown as mean□±□s.d. *p<0.05; **p<0.01; ***p<0.001; ****p<0.0001; NS, not significant.

**Extended Data Fig.3.**
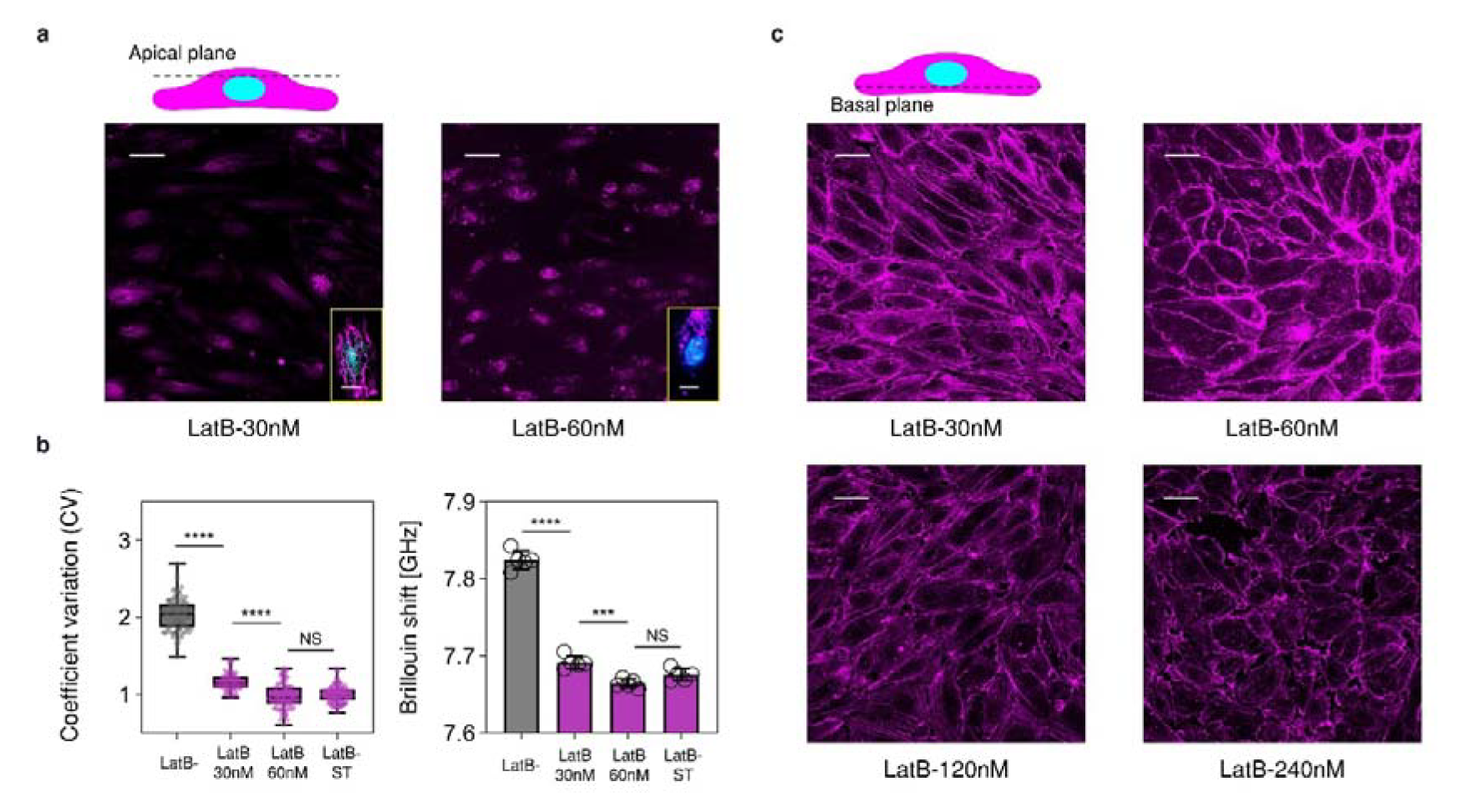
The effects of latrunculin B treatment on actin cap and basal conventional fibers structure. **a**, Effects of latrunculin B in various doses ranging from 30 to 60nM on actin cap formation. **b**, Comparison of LatB untreated, 30nM LatB treated, and 60nM treated cells under 24 hours USS, and the static control in coefficient variation (3 independent experiments, n > 80 cells) and Brillouin shift (5 independent experiments in Brillouin test, mean□±□s.d. *p<0.05; **p<0.01; ***p< 0.001; ****p<0.0001; NS, not significant). c, Effects of latrunculin B in various doses ranging from 30 to 240nM on basal conventional fibers. Scale bars, 40μm.

**Extended Data Fig.4.**
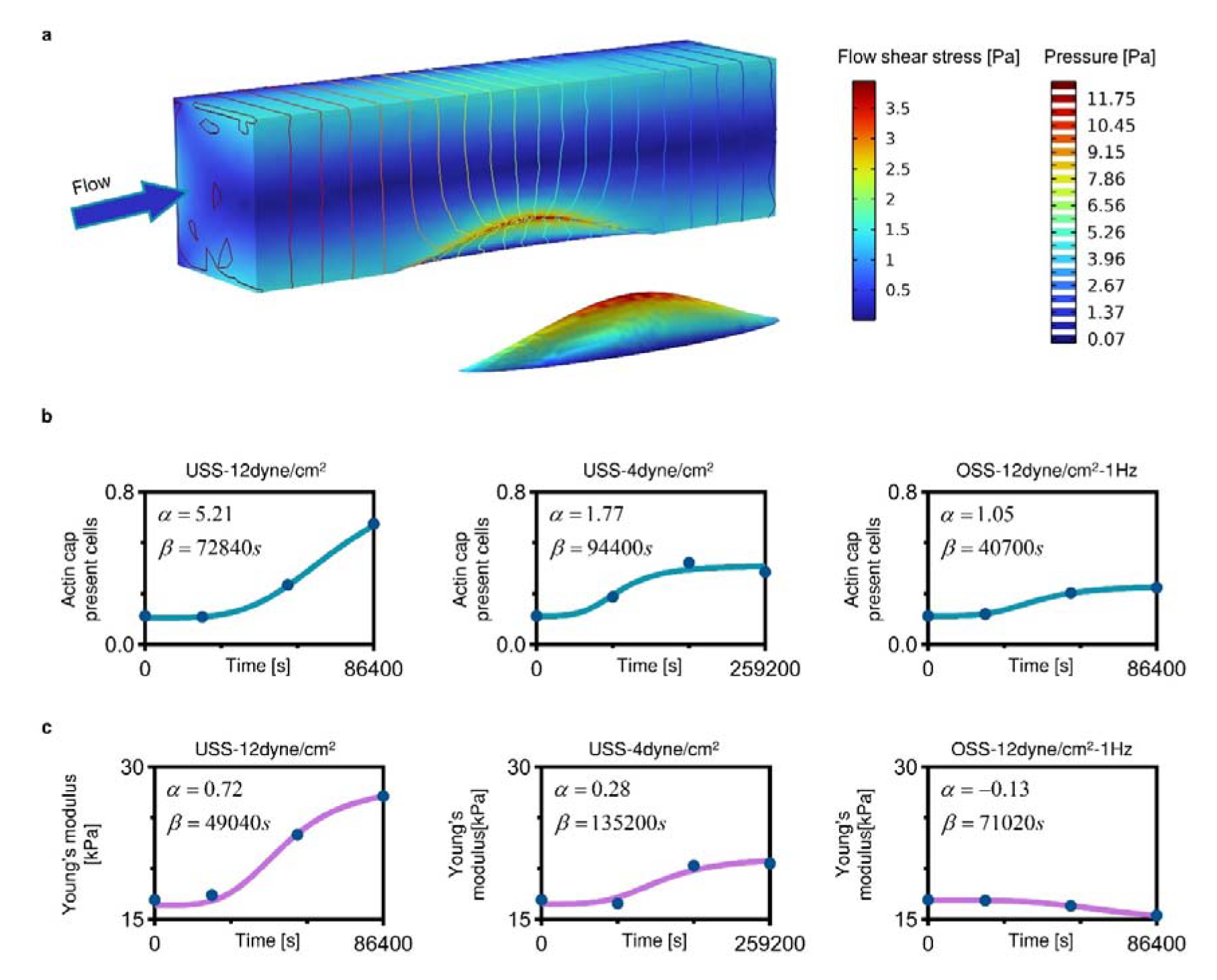
The fluid environment exerting on cell and experimental data-driven fitting. **a**, The diagram of shear stress at 12 dyne/cm^2^ and the pressure contour exerting on the cell membrane. **b,c**, The relationships between actin cap present cells nuclear Young’s modulus, flow time and shear stress condition are fitted from the experimental measurements with Hill equation (*H* = *H*_0_[1+ *αt*^4^/(*t*^4^ + *β*^4^)], α and β describe the shape of the Hill equation, which are shown in the corresponding subgraphs, the R-square value of each group fit exceeds 0.95).

**Extended Data Fig.5.**
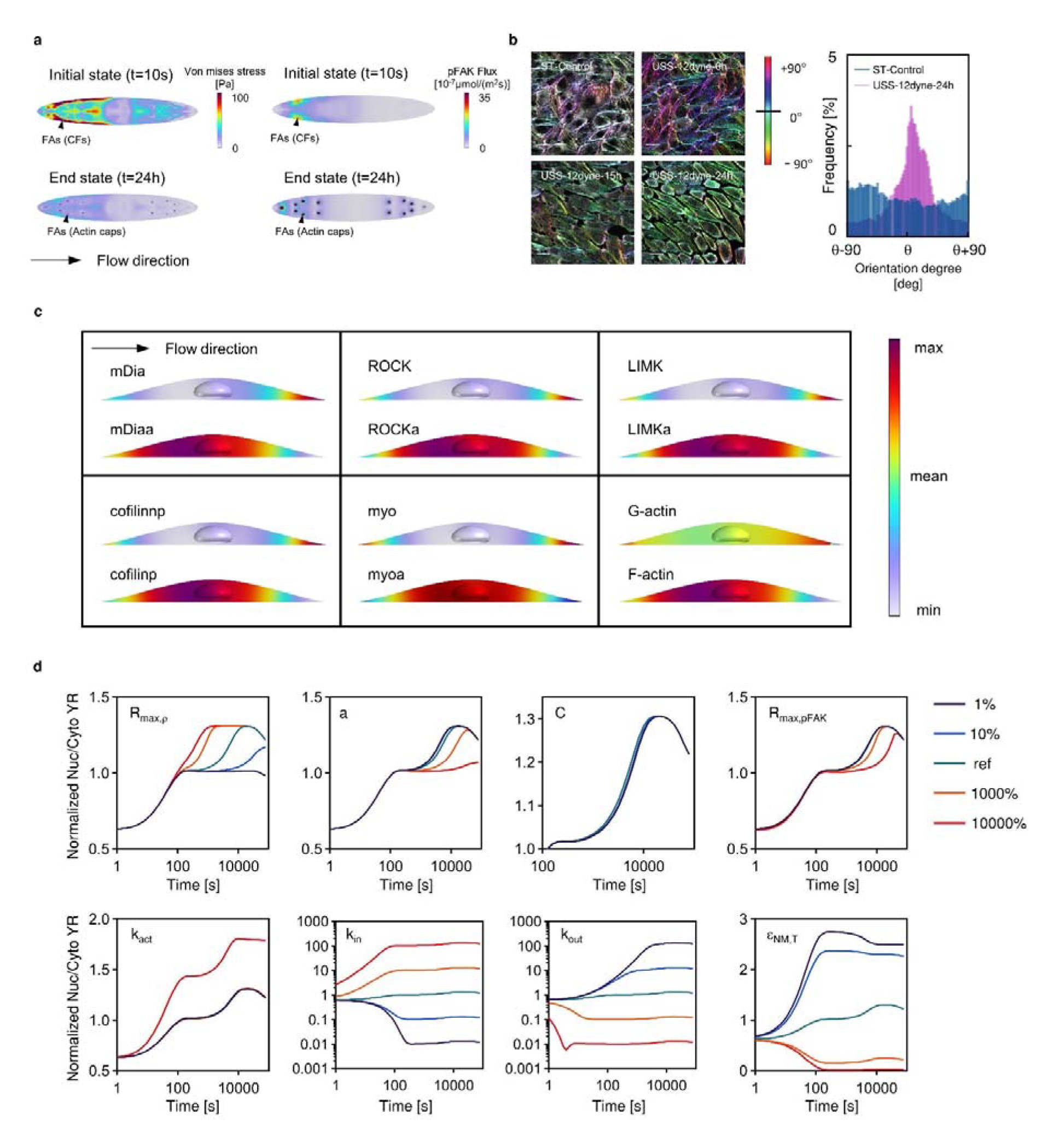
Signaling pathways in flow shear stress mechanochemical transduction process and the sensitivity analysis. **a,** The pFAK activation and the von mises stress in focal adhesion, which activates FAK. b, The representative conventional fibers polarization of endothelial cells under USS of 12 dyne/cm^2^. c, Signaling pathways downstream of RhoA, including the mDia, ROCK, LIMK, Cofilin, myosin, and actin. d, sensitivity analysis of key parameters involved in the mechanical process. Our analysis revealed that changes in the maximum activation rate of RhoA due to shear stress (R*_max,ρ)_*, the shear stress value when the activation rate of RhoA reaches *R_max,ρ_*/2 (*a*), the threshold von Mises stress at focal adhesion (*C*), the FAK phosphorylation rate (Rmax,pFAK), and the activation stress due to myosin activation (*k_act_*) do not significantly affect the biphasic translocation of YAP under unidirectional shear stress. However, alterations in the YAP nuclear import rate (*k_in_*), the YAP nuclear export rate (*k_out_*), and the threshold nuclear membrane strain of YAP nuclear import (*e*) – which are directly related to the nuclear-cytoplasmic properties of YAP – have a more substantial impact on the YAP transport.

**Extended Data Fig.6.**
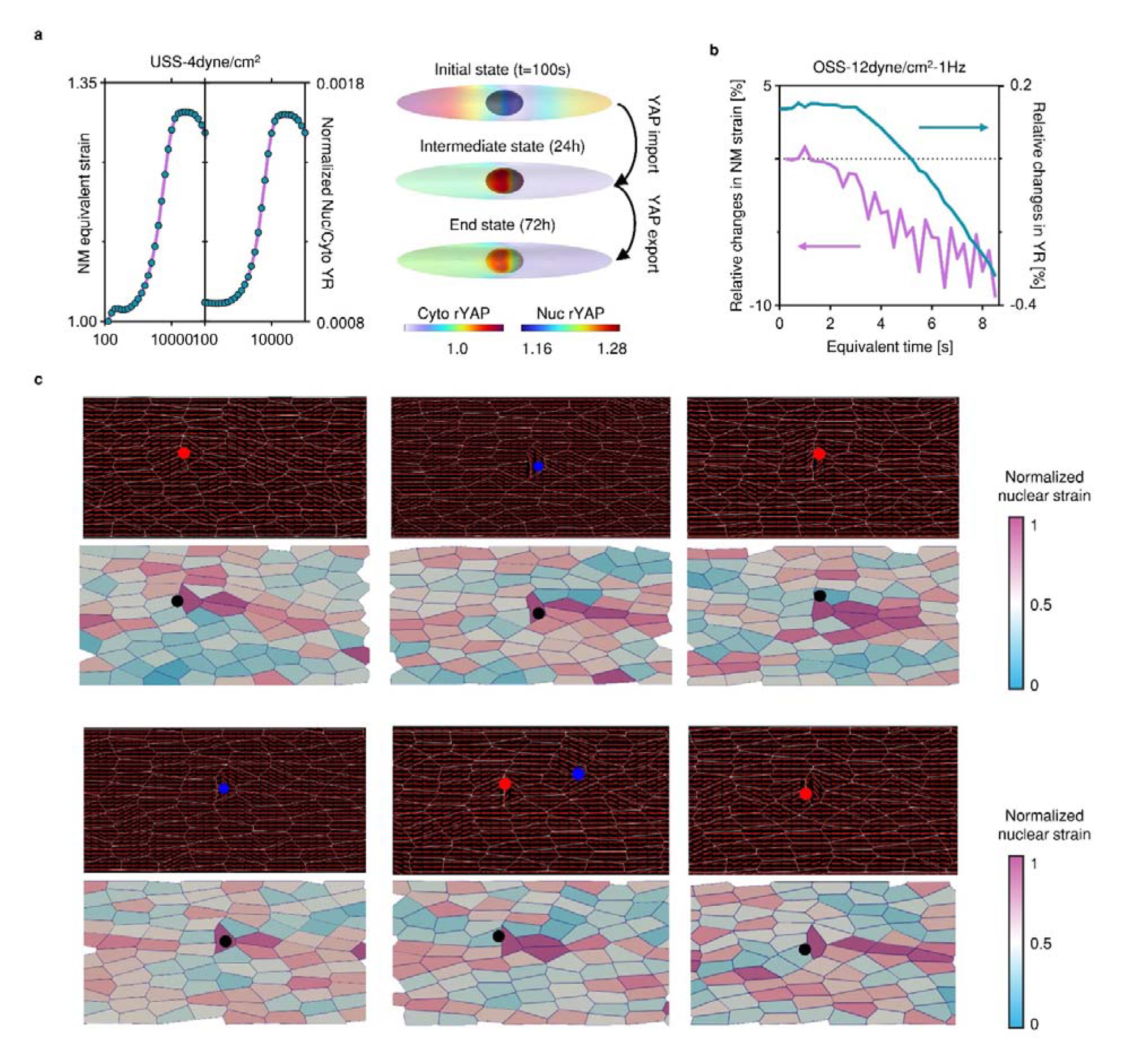
Mechanochemical modeling under 4 dyne/cm^2^ unidirectional shear stress and the topological defects formation induced by localized inhibiting the actin cap formation. **a,** The calculated nuclear membrane (NM) equivalent strain and YR changes in USS at 4 dyne/cm^2^ for 72 hours. **b,** the relative changes in NM strain and YR in OSS at 0±12 dyne/cm^2^, 1Hz by enhancing actin cap formation and inhibiting nuclear softening. **c,** The schematic of inhibition of actin cap in individual cells and its potential role in inducing topological defects and spatially heterogeneous distribution of nuclear membrane strain (fully inhibit the actin cap formation in single cell, red dots represent +1/2 defects, blue dots represent −1/2 defects).

