## Supplemental infomation for "Flow-induced mechanical coupling between perinuclear actin cap and nucleus governs spatiotemporal regulation of YAP transport"

**Supplementary Notes**

Tetrahedral mesh was used to mesh the entire geometric model. To ensure that grid nodes are aligned and non-overlapping, several subdomains with varying spatial sizes and transport properties shared topology. In the boundary layer, the element adjacent to the endothelial membrane surface was set to be 0.15μm, and gradually grew over 5 layers. As the boundary conditions for the fluid parts, the inlet velocity of the microfluidic channel was set up consistent with the experimental setup to ensure the shear stress magnitude of 4 dyne/cm^2^, 12 dyne/cm^2^, and 0 ± 12 dyne/cm^2^ (Sine function with the frequency of 1 Hz). The outlet pressure of the microfluidic channel was set to be a zero pressure. As the boundary conditions for the solid parts, the bottom of the compartment model was fixed in all directions. The surfaces of nucleus and actin cap are in contact with each other. The contact conditions were assumed as tied-type contact methods to mimics the present of LINC. The Fluid-Structure Interaction (FSI) interface combines the extracellular fluid domain with solid cytoplasm domain to capture the shear stress transmission. The FSI interface uses an arbitrary Lagrangian-Eulerian (ALE) method to combine fluid flow, represented using the Eulerian description and a spatial coordinate system, with solid mechanics, represented using the Lagrangian description and a material coordinate system. The boundary condition of the biochemical species transport was set as the flux condition to mimics the biochemical reaction at the (sub)cellular boundaries, including the basal plane, focal adhesion, nuclear membrane, and cell membrane. The unidirectional laminar flow simulations are performed for 24 or 72 hours in 0.1s timesteps, which grow exponentially to speed up the calculations. Oscillatory flow simulations shortened the process of actin cap formation and nucleus stiffness change over 24 hours, calculated in uniform 0.25s time steps for 8.64s to speed up the calculation.

**Supplementary Tables**

**Table S1 Mathematical equations for the components we considered in the computational model.**

| # | **Laminar flow** | |
| --- | --- | --- |
| 1 | 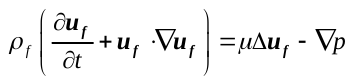  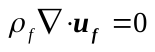 | |
|  | **Fluid solid interaction (FSI) interface** | |
| 1 | 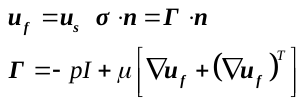 | |
|  | **Solid mechanics** | |
| 1 | General form 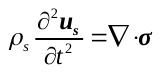 | |
| 2 | Cytoplasm  (linear elastic, isotropic) | 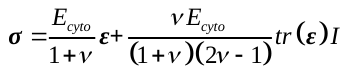 |
| 3 | Stress fiber  (strain hardening, along the direction of each stress fiber) | 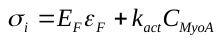  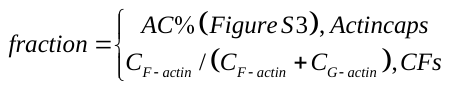  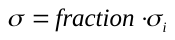 |
| 4 | Nucleus  (Maxwell model, isotropic) | 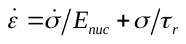 |
|  | **Diluted species transport** | |
|  | **Module 1: Flow Shear Stress Sensing** | |
|  | Equations | Reaction Rate |
| 1 | 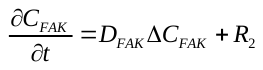  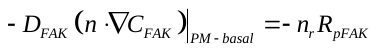 | 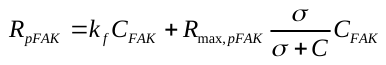  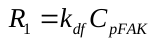 |
| 2 | 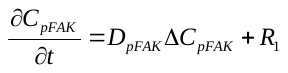  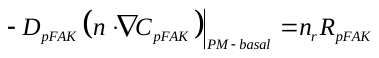 |  |
| 3 | 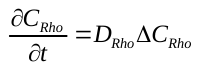  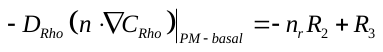  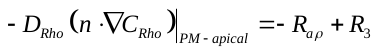 | 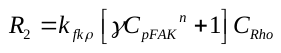  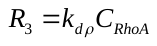  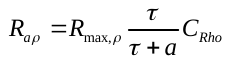 |
| 4 | 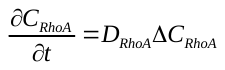  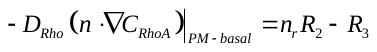  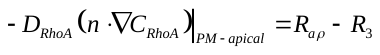 |  |
|  | **Module 2: Cytoskeleton regulation** | |
| 5 | 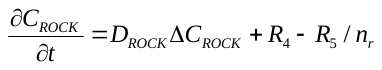 | 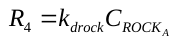  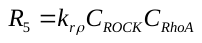 |
| 6 | 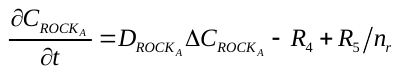 |  |
| 7 | 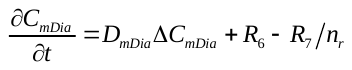 | 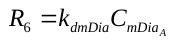  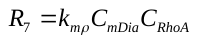 |
| 8 | 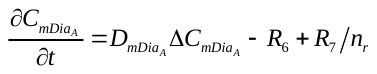 |  |
| 9 | 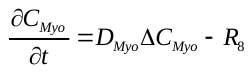 | 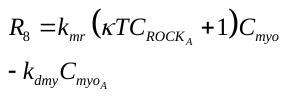 |
| 10 | 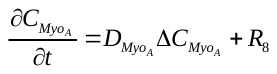 |  |
| 11 | 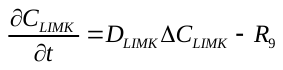 | 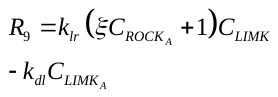 |
| 12 | 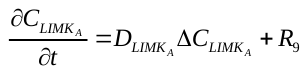 |  |
| 13 | 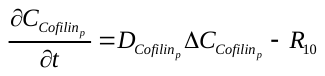 | 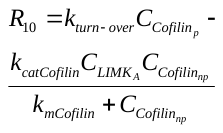 |
| 14 | 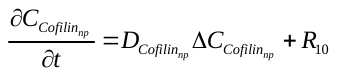 |  |
| 15 | 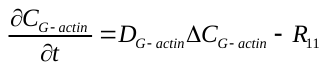 | 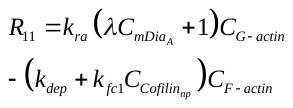 |
| 16 | 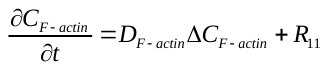 |  |
|  | **Module 3: YAP nucleocytoplasmic transport** | |
| 17 | 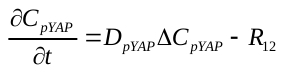 | 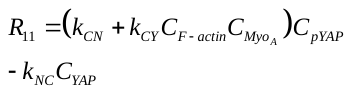  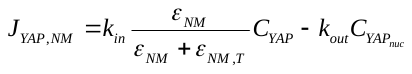 |
| 18 | 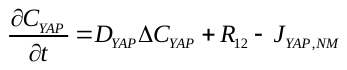 |  |
| 19 | 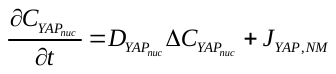 |  |

**Table S2 Parameters in computational model.**

| **Parameter** | **Value** | **Description** |
| --- | --- | --- |
| *ρ_f_* | 1000kg/m^3^ | Density of the fluid |
| *μ* | 0.95×10^-3^pa·s | Viscosity of the fluid |
| *ρ_cyto_* | 1060kg/m^3^ | Density of the cytoplasm |
| *E_cyto_* | 2.9kPa | Young’s modulus of the cytoplasm^1^ |
| *ρ_nuc_* | 1350kg/m^3^ | Density of the nucleus, estimated from^3^ |
| *E_nuc_* | Fitted from Brillouin measurement (see Extended Data Fig. 4c) | Young’s modulus of the nucleus, which is a function of flow time in each shear stress condition |
| *τ_r_* | 0.3s | Relaxation time of the nucleus^4,5^ |
| *ν* | 0.45 | Poisson ratio of the nucleus and cytoplasm^6^ |
| *R_max,pFAK_* | 0.379s^-1^ | FAK phosphorylation rate due to stress at focal adhesion applied by actin filaments^7^ |
| *k_f_* | 0.015s^-1^ | Baseline phosphorylation rate of FAK^7^ |
| *k_df_* | 0.035s^-1^ | Deactivation rate of FAK^8,9^ |
| *C* | 110Pa | Threshold von mises stress at focal adhesion, estimated based on^10^ |
| *k_fkρ_* | 1s^-1^ | Activation rate of RhoA at focal adhesion due to pFAK^9^ |
| *k_dρ_* | 0.625s^-1^ | Deactivation rate of RhoA^11^ |
| *γ* | 1000μM^-5^ | Activation rate of RhoA due to pFAK^12^ |
| *n* | 5 | Cooperativity between FAK and Rho^12,13^ |
| *R_aρ_* | 1.32s^-1^ | Maximum activation rate of RhoA due to the shear stress, which is selected to close the order of magnitude for RhoA activation rate due to pFAK^12^ |
| *a* | 0.5Pa | Threshold shear stress at cell membrane^14^ |
| *k_drock_* | 0.8s^-1^ | Deactivation rate of ROCK^9,15^ |
| *k_rρ_* | 0.648s^-1^μM^-1^ | Activation rate of ROCK due to RhoA^16^ |
| *k_dmdia_* | 0.005s^-1^ | Deactivation rate of mDia^17^ |
| *k_mr_* | 0.03s^-1^ | Activation rate of mDia due to RhoA^16^ |
| *κ* | 36μM^-1^ | Amplification of myosin to ROCK^16^ |
| *k_dmy_* | 0.067s^-1^ | Deactivation rate of activated myosin^9,12^ |
| *K_F_* | 1.45MPa | Young’s modulus of stress fiber^2^ |
| *k_act_* | 1000J/mol  **σ_i_=K_F_ε+σ_M_, σ_M_=k_act_C_MyoA_* | Activate stress due to myosin activation, estimated by matching the magnitude order of passive stress in actin fibers |
| *k_lr_* | 0.07s^-1^ | Activation rate of LIMK^9,12^ |
| *ξ* | 55.49μM^-1^ | Amplification of LIMK to ROCK^12^ |
| *k_dl_* | 2s^-1^ | Deactivation rate of LIMK_A_^9,12^ |
| *k_turn-over_* | 0.04s^-1^ | Activation rate of cofilin^9,12^ |
| *k_catCofilin_* | 0.34s^-1^ | Rate of catalysis for cofilin phosphorylation^18^ |
| *k_mcofilin_* | 4μM | Concentration of cofilin when the reaction rate reaches its half of max rate^18^ |
| *k_ra_* | 0.4s^-1^ | Activation (polymerization) rate of F-actin^9,12^ |
| *λ* | 50μM^-1^ | Amplification of F-actin to mDia^9,12^ |
| *k_dep_* | 3.5s^-1^ | Deactivation (depolymerization) rate of F-actin^9,12^ |
| *k_fc1_* | 4s^-1^μM^-1^ | Deactivation (depolymerization) rate of F-actin due to cofilin^9,12^ |
| *k_CN_* | 0.56s^-1^ | Baseline rate of YAP dephosphorylation^12,19,20^ |
| *k_CY_* | 7.6×10^-4^μM^-1^s^-1^ | Rate of YAP dephosphorylation due to stress fiber^12,19,20^ |
| *ε_NM,T_* | 0.0002 | Threshold first principal strain of YAP nuclear import, estimated by matching the magnitude order of the calculated strain in pure mechanical model |
| *k_NC_* | 0.14s^-1^ | Rate of YAP phosphorylation^12,19,20^ |
| *k_in_* | 10s^-1^μM^-1^ | Maximum rate of YAP nuclear import^12,19,20^ |
| *k_out_* | 10s^-1^μM^-1^ | Nuclear export rate of YAP^21^ |
